# ATP-independent phosphate recycling on AGC kinase activation loops induced by alkali metal ions

**DOI:** 10.1101/2025.09.04.674365

**Authors:** Koji Kubouchi, Hideyuki Mukai

## Abstract

Changes in extracellular Na⁺ and K⁺ concentrations have traditionally been considered to influence intracellular signal transduction through alterations in cell volume or membrane potential. However, whether intracellular ion concentration changes directly regulate signaling molecules, independent of these conventional pathways, remains largely unexplored. In this study, we demonstrate that even in the absence of cellular membranes, an increase in Na⁺ or K⁺ concentration rapidly reduces activation-loop phosphorylation of multiple AGC kinases, including PKN, PKCζ/λ, and p70 S6 kinase. When ion concentrations were reduced, the activation-loop phosphorylation, which had initially decreased, recovered within a short period. Notably, this recovery occurred in the absence of PDK1, a known kinase responsible for the phosphorylation of these activation loops, and did not require ATP or Mg²⁺ in lysate assays. ³²P tracing experiments revealed a novel ‘reacquisition of phosphate group’ mechanism, in which phosphate groups transiently dissociate from the activation loops under high Na⁺ or K⁺ conditions and are subsequently re-incorporated into the activation loops when ion concentrations are reduced. These findings indicate that elevated Na⁺ or K⁺ concentrations directly and rapidly reduce the activity of multiple AGC kinases, and that the activity rapidly recovers upon reduction in ion concentrations, through an unconventional phosphate transfer mechanism distinct from canonical protein phosphorylation reactions. Our study suggests the existence of a robust phosphorylation homeostasis mechanism independent of conventional kinase-phosphatase systems, providing new insights into signaling pathways regulated by intracellular Na⁺/K⁺ ion dynamics.

## Introduction

Sodium (Na⁺) and potassium (K⁺) are essential ions in biological systems and play central roles in a wide range of physiological functions. These ions regulate various processes, including the osmotic gradient between intracellular and extracellular environments, membrane potential (Pivovarov et al., 2018; Yu & Catterall, 2003), maintenance of cell volume (Hoffmann et al., 2009; Strange, 2004), intracellular pH control (Doyen et al., 2022), and glucose uptake (Thorsen et al., 2014). Additionally, they are crucial for neuronal action potential generation and propagation (Pivovarov et al., 2018; Yu & Catterall, 2003), muscle contraction (McKenna et al., 2024), fluid and electrolyte balance in the kidneys (John, 2006), and hormone secretion (Ghatta et al., 2006). Therefore, abnormalities in Na⁺ and K⁺ homeostasis are directly linked to various pathological conditions, including hypertension (Adrogué & Madias, 2007), cardiac arrhythmias (Gary, 2016), and muscle dysfunction (Goldin, 2001). It is widely recognized that changes in Na⁺ and K⁺ concentrations influence intracellular signal transduction through alterations in cell volume and membrane potential. Specifically, changes in cell volume can activate signaling molecules such as mitogen-activated protein kinases (MAPK) and phosphoinositide 3-kinase (PI3K)/AKT pathways via osmotic regulation and mechanical stress responses (Hoffmann et al., 2009; Mongin & Orlov, 2001). Meanwhile, alterations in membrane potential regulate calcium signaling via voltage-dependent ion channels, impacting transcription factor activation and metabolic responses (Berridge, 1998, 2008). However, the possibility that changes in Na⁺ and K⁺ concentrations directly modulate intracellular signaling independent of cell volume or membrane potential changes remains largely unexplored.

Protein phosphorylation plays a crucial role in intracellular signal transduction, influencing protein-protein interactions, subcellular localization, stability, and enzymatic activity (Ardito et al., 2017; Nishi et al., 2011; Pang et al., 2022). Phosphorylation events often function as molecular switches that regulate downstream signaling pathways. This process is catalyzed by protein kinases, which transfer the γ-phosphate group from ATP to specific amino acid residues in target proteins (Adams, 2001; Roskoski, 2015). In mammalian cells, serine (Ser), threonine (Thr), and tyrosine (Tyr) residues are the primary phosphorylation targets. The mammalian kinome, comprising structurally conserved catalytic domains, is categorized into eight major kinase families: AGC, CAMK, CK1, CMGC, STE, TK, TKL, and Others (Manning et al., 2002). Among these, phosphorylation of the activation loop within the catalytic domain is essential for the enzymatic activity of kinases belonging to the CMGC (Kannan & Neuwald, 2004), AGC (Leroux et al., 2018), and TK families (Hubbard & Till, 2000). The AGC family includes kinases such as protein kinase N (PKN), protein kinase C (PKC), p70 S6 kinase (p70S6K/RPS6KB1), AKT (protein kinase B/PKB), and serum- and glucocorticoid-induced kinase (SGK), which are involved in cell proliferation, protein synthesis, cell cycle regulation, intracellular transport, and cell motility (Bahrami et al., 2014; Dempsey et al., 2000; Franke, 2008; Lang et al., 2006; Mukai, 2003; Sophocleous et al., 2021). Furthermore, dysregulation of AGC kinases is implicated in various diseases, including cancer, diabetes, immune disorders, and neurodegenerative diseases (Hers et al., 2011; Koivunen et al., 2006; Leenders et al., 2004; Manning & Toker, 2017; Mochly-Rosen et al., 2012; Peifer & Alessi, 2008; Sophocleous et al., 2021; Standaert et al., 1999; Yeo et al., 2017). Regulation of activation-loop phosphorylation in AGC kinases is, therefore, a critical research topic for understanding disease mechanisms and developing therapeutic strategies.

The 3-phosphoinositide-dependent protein kinase 1 (PDK1) is the responsible kinase for activation-loop phosphorylation of AGC family kinases, including AKT (PKB) and p70 S6 kinase, and is known as a “master kinase” due to its central role in signal transduction (Leroux et al., 2018; Mora et al., 2004; Zheng et al., 2023). Although more than a quarter of a century has passed since the discovery of PDK1-mediated phosphorylation of AKT (Alessi et al., 1997), studies on PDK1-independent activation-loop phosphorylation mechanisms remain largely unknown. Elucidating potential PDK1-independent regulatory mechanisms of activation-loop phosphorylation could provide new insights into intracellular signaling and novel therapeutic strategies.

In this study, we investigated the possibility that changes in Na⁺ and K⁺ concentrations modulate intracellular signaling independently of cell volume or membrane potential changes. We found that treatment of various animal cell lines with high concentrations of Na⁺ or K⁺ led to a rapid and marked decrease in activation-loop phosphorylation of multiple AGC kinases, even in the absence of cellular membranes. Notably, the activation-loop phosphorylation rapidly recovered upon subsequent reduction of ion concentrations. Importantly, this recovery occurred independently of ATP and Mg²⁺, and the dissociated phosphate group was selectively re-incorporated into the activation loop. These findings suggest that changes in Na⁺ and K⁺ concentrations can rapidly and potently regulate AGC kinase activity and that a previously unreported mechanism of phosphate group transfer underlies this regulation.

## Results

### Elevated extracellular NaCl concentrations rapidly reduce the PKN activation-loop phosphorylation in COS7 cells

To investigate whether changes in extracellular NaCl concentration affect the properties of intracellular PKN, we examined the activation-loop phosphorylation in PKN1, a critical site for its activity, in COS7 cells derived from the African green monkey kidney. Cells were incubated in media containing NaCl at concentrations of 300 mM, 450 mM, and 600 mM for 10 min, respectively, followed by cell lysis and immunoblotting using phospho-specific antibodies. The results demonstrated a clear decrease in the activation-loop phosphorylation in all PKN isoforms (PKN1, PKN2, and PKN3) upon incubation with 450 mM or 600 mM NaCl (Figure 1A).

**Figure 1.**
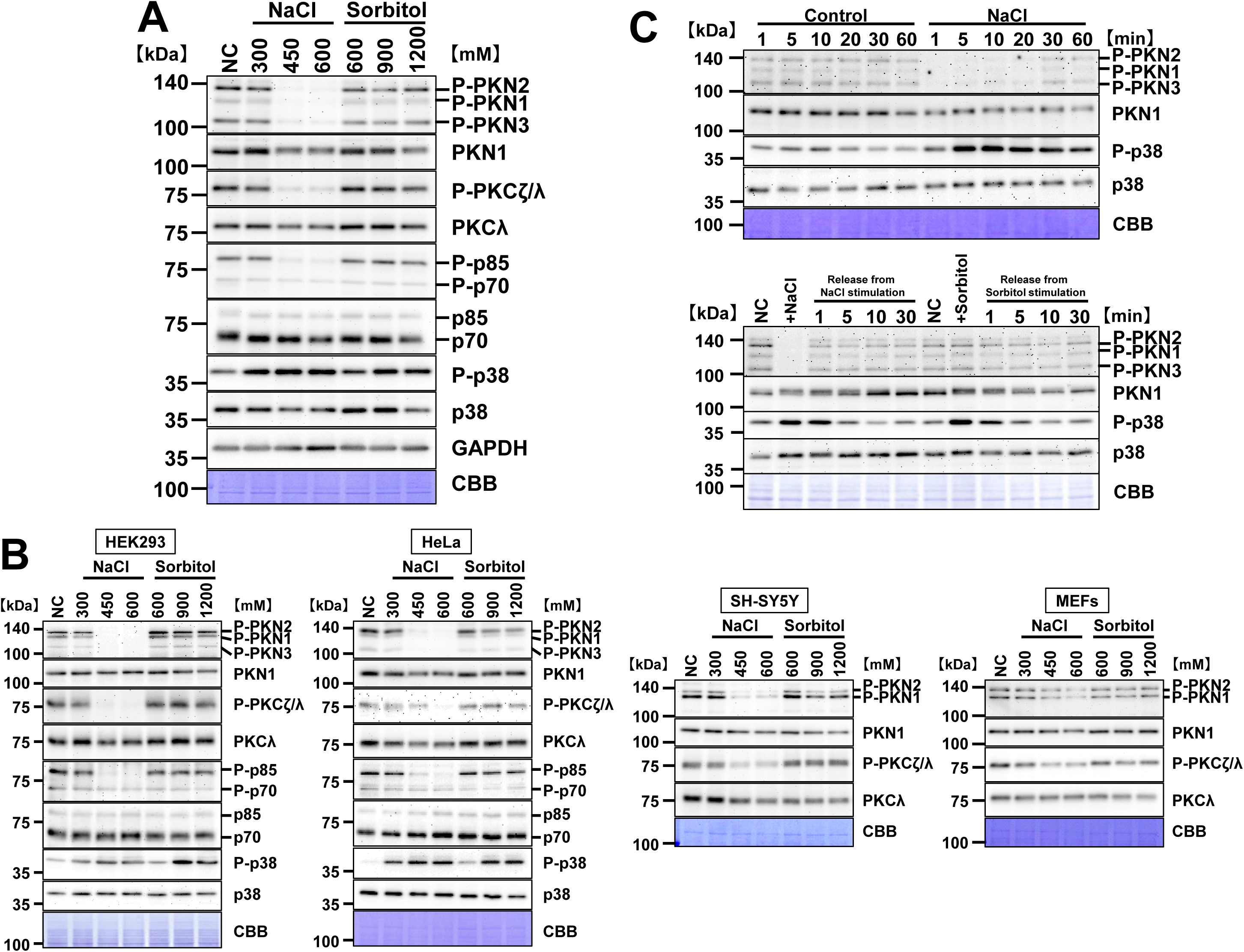
NaCl, but not isosmotic sorbitol, reversibly reduces AGC kinase activation-loop phosphorylation in diverse cultured cells. Cells were incubated for 10 min in DMEM supplemented with NaCl or sorbitol at the indicated concentrations (mM). Whole-cell lysates were analyzed by immunoblotting with phospho-specific antibodies to activation-loop sites together with total proteins where indicated; membranes were stained with Coomassie Brilliant Blue (CBB) after immunoblotting as a loading control. GAPDH was included as an additional loading control where specified. (A) Dose–response immunoblots showing NaCl-induced decrease in activation-loop phosphorylation in PKN (PKN1/2/3), atypical PKC (PKCζ/λ), and p70 S6 kinase (p70/p85), whereas sorbitol had little or no effect under the same osmolarities. Total PKCλ, total p70/p85, and GAPDH are shown together with CBB. (B) The NaCl-induced decrease in activation-loop phosphorylation in PKN and PKCζ/λ in multiple cell types, including HEK293, HeLa, SH-SY5Y, and primary wild-type mouse embryonic fibroblasts (MEFs). (C) Reversible decrease in phosphorylation levels at the PKN activation loop. COS7 cells were treated with 600 mM NaCl for the indicated durations (upper panel), or incubated for 10 min with 600 mM NaCl or 1200 mM sorbitol followed by washout into DMEM for the indicated recovery times (lower panels). NaCl caused a reversible decrease in activation-loop phosphorylation of PKN, whereas sorbitol did not. Phospho-p38 (P-p38) and total p38 are shown as markers of hypertonic-stress signaling. Representative immunoblots are shown. The following figure supplements are available for figure 1: **Figure supplement 1.** Effect of varying NaCl concentrations on activation-loop phosphorylation of PKN and PKCζ/λ in COS7 cells. **Figure supplement 2.** KCl-induced decrease in PKN phosphorylation in insect cell extracts (Sf9) but not in *E. coli* extracts.

Next, we examined other AGC family kinases that have a highly conserved activation-loop sequence with PKN, including PKCζ/λ and p70 S6 kinase (Belham et al., 1999; Parekh et al., 2000), using their respective phospho-specific antibodies. Immunoblotting revealed a clear decrease in activation-loop phosphorylation at 450 mM and 600 mM NaCl in both cases (Figure 1A).

### The effect of NaCl is independent of osmotic stress

To determine whether the observed effects were due to changes in osmotic pressure rather than NaCl itself, we supplemented the culture medium with sorbitol at concentrations that matched the osmolarity of the NaCl treatments. In this case, no decrease in the phosphorylation levels of any AGC family kinases was observed (Figure 1A). However, p38 MAPK, a member of the CMGC kinase family known to be activated by hyperosmotic stress (Ben Messaoud et al., 2015; Hoffmann et al., 2009), showed an increase in activation-loop phosphorylation in response to both NaCl and sorbitol treatments (Figure 1A), consistent with previous reports. These results indicate that the NaCl-induced decrease in AGC kinase phosphorylation occurs through a mechanism distinct from osmotic stress.

### The NaCl-induced decrease in activation-loop phosphorylation is not cell-type specific and is rapidly reversible

To assess whether the decrease in activation-loop phosphorylation was specific to COS7 cells, we performed similar experiments in other cell lines, including human embryonic kidney HEK293 cells, human cervical cancer HeLa cells, human neuroblastoma SH-SY5Y cells, and mouse embryonic fibroblasts (MEFs). Despite some variations in sensitivity among cell types, all tested cell lines exhibited a clear decrease in activation-loop phosphorylation upon high NaCl treatment (Figure 1B). This suggests that the NaCl-induced suppression of AGC kinase activity is a general phenomenon in mammalian cells.

Furthermore, we found that the decrease in activation-loop phosphorylation occurred within 1 min of NaCl exposure and was rapidly recovered to near-baseline levels within 1 min after returning the cells to normal medium (Figure 1C). In contrast, lowering the NaCl concentration below physiological levels did not alter PKN activation-loop phosphorylation (Figure 1 - figure supplement 1). These findings suggest that exposure to high NaCl concentrations leads to a rapid and marked suppression of multiple AGC kinases, and the subsequent normalization of NaCl concentration results in a swift recovery of kinase activities.

### The decrease in PKN activation-loop phosphorylation is induced by the addition of alkali metal ions (except Li^+^) and is observed even in a cell-free system

To determine whether the decrease in activation-loop phosphorylations in PKN, PKCζ/λ, and p70 S6 kinase was specific to high NaCl treatment, we examined the effects of substituting NaCl with other salts. COS7 cells were incubated for 10 min in media supplemented with LiCl, KCl, RbCl, or CsCl, followed by immunoblotting to assess phosphorylation levels. The results showed that the presence of KCl, RbCl, and CsCl except LiCl led to a marked decrease in PKN activation-loop phosphorylation, similar to the effects observed with NaCl treatment (Figure 2A). Furthermore, KCl, RbCl, and CsCl induced this effect at lower concentrations than NaCl, with KCl being the most effective in reducing phosphorylation levels. Additionally, when COS7 cells were incubated in media containing KHCO_3_ or KI, a similar decrease as in case of KCl in PKN activation-loop phosphorylation was observed (Figure 2B). These results suggest that high concentrations of alkali metal ions, including Na⁺, K⁺, Rb⁺, and Cs⁺, contribute to the decrease in PKN activation-loop phosphorylation.

**Figure 2.**
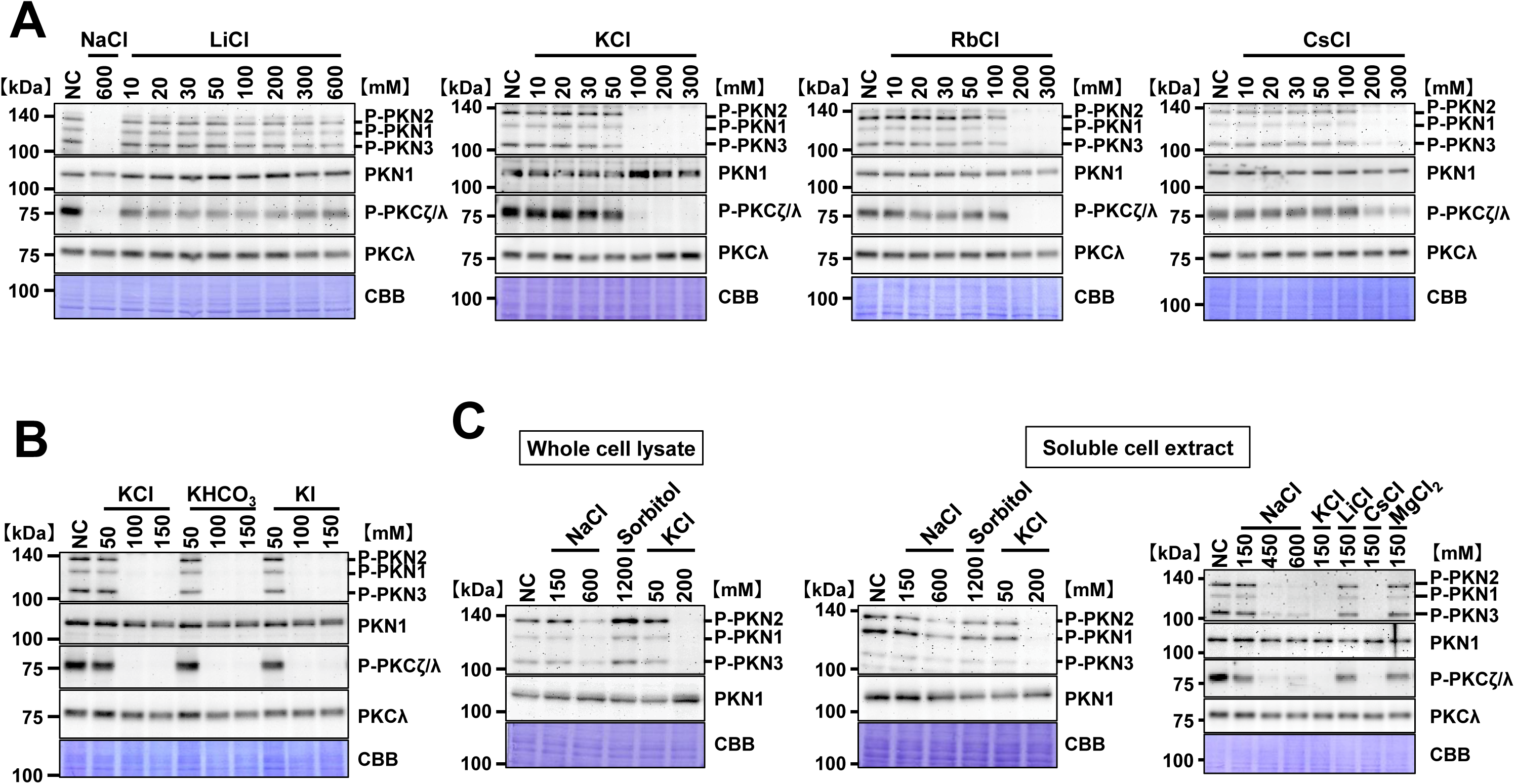
Alkali metal cations reduce activation-loop phosphorylation of PKN (and atypical PKC) and the effect is reproduced in cell lysates. COS7 cells (or lysates, as indicated) were incubated for 10 min with the indicated salts at the indicated concentrations (mM). Whole-cell lysates were analyzed by immunoblotting with phospho-specific antibodies against activation-loop sites together with the corresponding total proteins where shown. Membranes were stained with Coomassie Brilliant Blue (CBB) after immunoblotting as a loading control; PKCλ and PKN1 are included as total-protein references where indicated. Representative immunoblots are shown. (A) Effect of alkali metal ions on the phosphorylation levels in the activation loops of PKN and PKCζ/λ. KCl, RbCl, and CsCl reduced activation-loop phosphorylation of PKN (PKN1/2/3) and PKCζ/λ in intact COS7 cells in a concentration-dependent manner as well as NaCl. (B) Effect of K^+^ salts on the activation-loop phosphorylation of PKN and PKCζ/λ. K^+^ reduced activation-loop phosphorylation regardless of the accompanying anion. (C) Decrease in activation-loop phosphorylation of PKN and PKCζ/λ *in vitro*. Whole-cell lysates were incubated with NaCl, KCl, or sorbitol; soluble cell extracts were incubated with LiCl, NaCl, KCl, CsCl and MgCl₂, or sorbitol, all for 10 min at the indicated mM. The following figure supplement is available for figure 2: **Figure supplement 1.** Effects of membrane potential and osmotic pressure on PKN activation-loop phosphorylation.

Elevated extracellular Na⁺ and K⁺ concentrations are known to cause plasma membrane depolarization (Bortner et al., 2001; Davis et al., 2022), which in turn leads to Ca²⁺ influx and subsequent activation of intracellular Ca²⁺ signaling (Berridge, 1998; Cárdenas et al., 2004; Macías et al., 2001; Oliveria et al., 2007; Schmitt et al., 2004). To investigate whether the observed decrease in PKN activation-loop phosphorylation required Ca²⁺ influx, COS7 cells were incubated in Ca²⁺-free medium for 10 min, followed by treatment with high concentrations of NaCl or KCl. The results demonstrated that the decrease in PKN activation-loop phosphorylation occurred even in the absence of extracellular Ca²⁺ (Figure 2 - figure supplement 1A). Furthermore, to examine whether changes in osmolarity were required for this decrease, we exchanged the extracellular concentrations of NaCl and KCl while maintaining osmolarity, and found that the decrease in PKN activation-loop phosphorylation was still induced (Figure 2 - figure supplement 1B).

Based on these findings, we hypothesized that the decrease in PKN activation-loop phosphorylation is independent of changes in membrane potential or osmotic pressure. To further explore this possibility, we examined the effects of high Na⁺ and K⁺ concentrations in a cell-free system. Whole-cell lysates and soluble cell extracts were prepared from COS7 cells through ultrasonic disruption and centrifugation at 20,400 × g for 30 min. These fractions were then supplemented with high concentrations of NaCl or KCl, followed by immunoblotting to assess phosphorylation levels (Figure 2C). The results showed that, whether using whole-cell lysates or soluble cell extracts, high Na⁺ or K⁺ concentrations led to a decrease in PKN activation-loop phosphorylation. These findings indicate that the reversible changes in AGC kinase activation-loop phosphorylation can occur not only in intact cells but also in a cell-free soluble extract upon direct addition of Na⁺ or K⁺.

### Protein phosphatase 2A (PP2A) is involved in the Na⁺- or K⁺-induced decrease in PKN activation-loop phosphorylation

Our previous studies demonstrated that preincubation of GST-tagged PKN1 purified from Sf9 cells with protein phosphatase 2A (PP2A) reduces the kinase activity of PKN1 (Yoshinaga et al., 1999). Additionally, we found that PKN1 associates with the scaffolding protein CG-NAP (also known as AKAP450), which serves as an anchoring adapter for both phosphatase 1 (PP1) and PP2A (Takahashi et al., 1999). Based on these findings, we investigated whether PP1 or PP2A contributes to the Na⁺- and K⁺-induced decrease in PKN activation-loop phosphorylation. First, we treated COS7 cells with okadaic acid (Cohen et al., 1990), a potent inhibitor of PP1 and PP2A, followed by incubation in NaCl-supplemented medium. Immunoblot analysis revealed that okadaic acid treatment significantly suppressed the NaCl-induced decrease in PKN and PKC activation-loop phosphorylations (Figure 3A). Next, we assessed the effects of rubratoxin A (Wada et al., 2010), a selective inhibitor of PP2A, in soluble extracts of COS7 cells on the activation-loop phosphorylation. As shown in Figure 3B, rubratoxin A inhibited the decrease in PKN and PKC activation-loop phosphorylation upon Na⁺ or K⁺ (Figure 3B). Furthermore, when PP2A catalytic subunit was knocked down in HeLa cells using siRNA, the NaCl- or KCl-induced decrease in PKN activation-loop phosphorylation was suppressed (Figure 3C). These results suggest that PP2A plays a critical role in mediating the decrease in PKN activation-loop phosphorylation upon Na⁺ or K⁺ exposure. Previous reports have indicated that PP2A activity is influenced by metal ions such as Mn²⁺ and Co²⁺ (Cai et al., 1995). To examine whether Na⁺ or K⁺ directly activates PP2A, we purified recombinant PP2A complexes from Sf9 cells and performed *in vitro* phosphatase assays in the presence or absence of NaCl or KCl. The results demonstrated that neither NaCl nor KCl enhanced PP2A phosphatase activity; instead, both salts inhibited its activity (Figure 3D). PP2A is required for the Na^+^ or K^+^ -induced decrease in activation-loop phosphorylation, but PP2A itself is unlikely to be directly activated by these salts *in vitro*.

**Figure 3.**
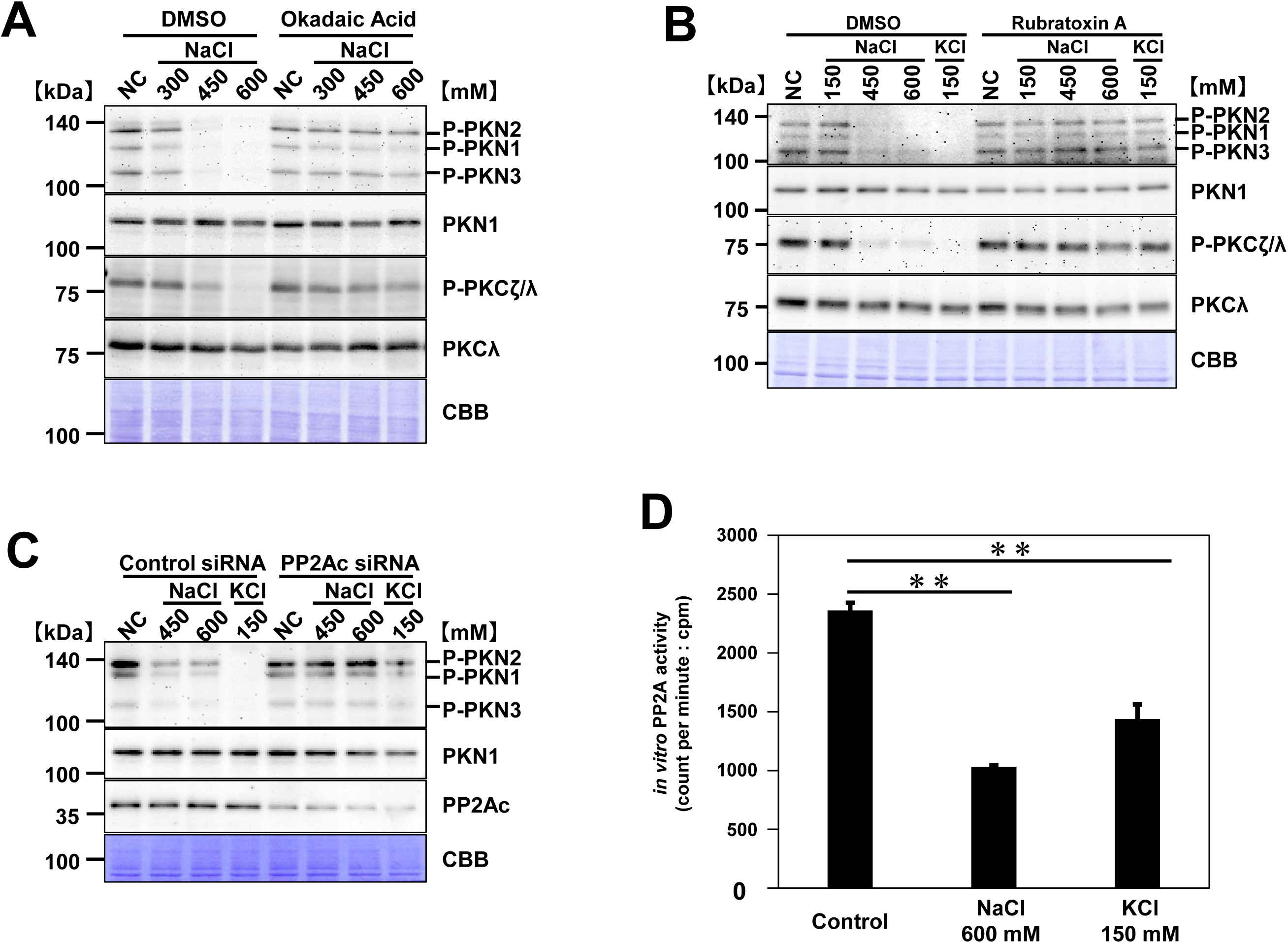
Protein phosphatase PP2A mediates the decrease in PKN activation-loop phosphorylation induced by NaCl or KCl. COS7 (or HeLa, as indicated) cells or lysates were treated with salts for 10 min at the indicated concentrations (mM). Whole-cell lysates were analyzed by immunoblotting with phospho-specific antibodies to activation-loop sites of PKN (PKN1/2/3) and PKCζ/λ, together with the corresponding total proteins where indicated. Membranes were stained with Coomassie Brilliant Blue (CBB) as a loading control. Representative immunoblots are shown. (A) Effect of okadaic acid on the decrease in activation-loop phosphorylation. COS7 cells were pre-incubated with DMSO or 250 nM okadaic acid for 3 h at 37°C (5% CO_2_). The cells were then transferred to DMEM containing NaCl at the indicated concentrations while okadaic acid (or DMSO) was kept at the same concentration during the entire 10 min NaCl treatment. (B) Effect of rubratoxin A on the decrease in activation-loop phosphorylation. COS7 soluble extracts were pre-treated with DMSO or 3 µM rubratoxin A for 10 min 30°C, after which they were incubated for an additional 10 min with NaCl or KCl at the indicated concentrations in the continued presence of rubratoxin A (or DMSO). (C) Effect of PP2A knockdown on the activation-loop phosphorylation. HeLa cells were transfected with 10 nM control siRNA or siRNA targeting the catalytic subunit of PP2A, and then cultured for 48 h at 37°C (5% CO_2_). After incubation, these cells were treated with DMEM supplemented with NaCl or KCl at the indicated concentrations for 10 min. Immunoblot analysis of the HeLa cell lysates was performed using antibodies against the total and phosphorylated forms of PKN and PP2A catalytic subunit. (D) Effects of NaCl and KCl on PP2A phosphatase activity *in vitro*. PP2A phosphatase activity was measured using radiolabeled phospho-casein as a substrate. Data are mean ± SEM. Data were analyzed using an unpaired t-test (n = 3). **p < 0.01. The input radioactivity of the phospho-casein was 21,324.49 cpm.

### Recovery of PKN activation-loop phosphorylation after normalization of K⁺ concentration is independent of PDK1 and does not require ATP or Mg²⁺

To investigate the mechanism underlying the rapid recovery of activation-loop phosphorylation following its initial decrease induced by elevated Na⁺ or K⁺ concentrations, we focused primarily on K⁺, which exerted stronger effects at lower concentrations.

We first tested whether the recovery of activation-loop phosphorylation in PKN, PKCζ/λ, and p70 S6 kinase depends on phosphoinositide-dependent protein kinase 1 (PDK1), a well-established upstream kinase for these AGC kinases. Mouse embryonic fibroblasts (MEFs) were treated with high concentrations of KCl in the presence or absence of wortmannin, a PI3K inhibitor that acts upstream of PDK1. Immunoblot analysis showed that wortmannin did not affect the recovery of activation-loop phosphorylation (Figure 4A).

**Figure 4.**
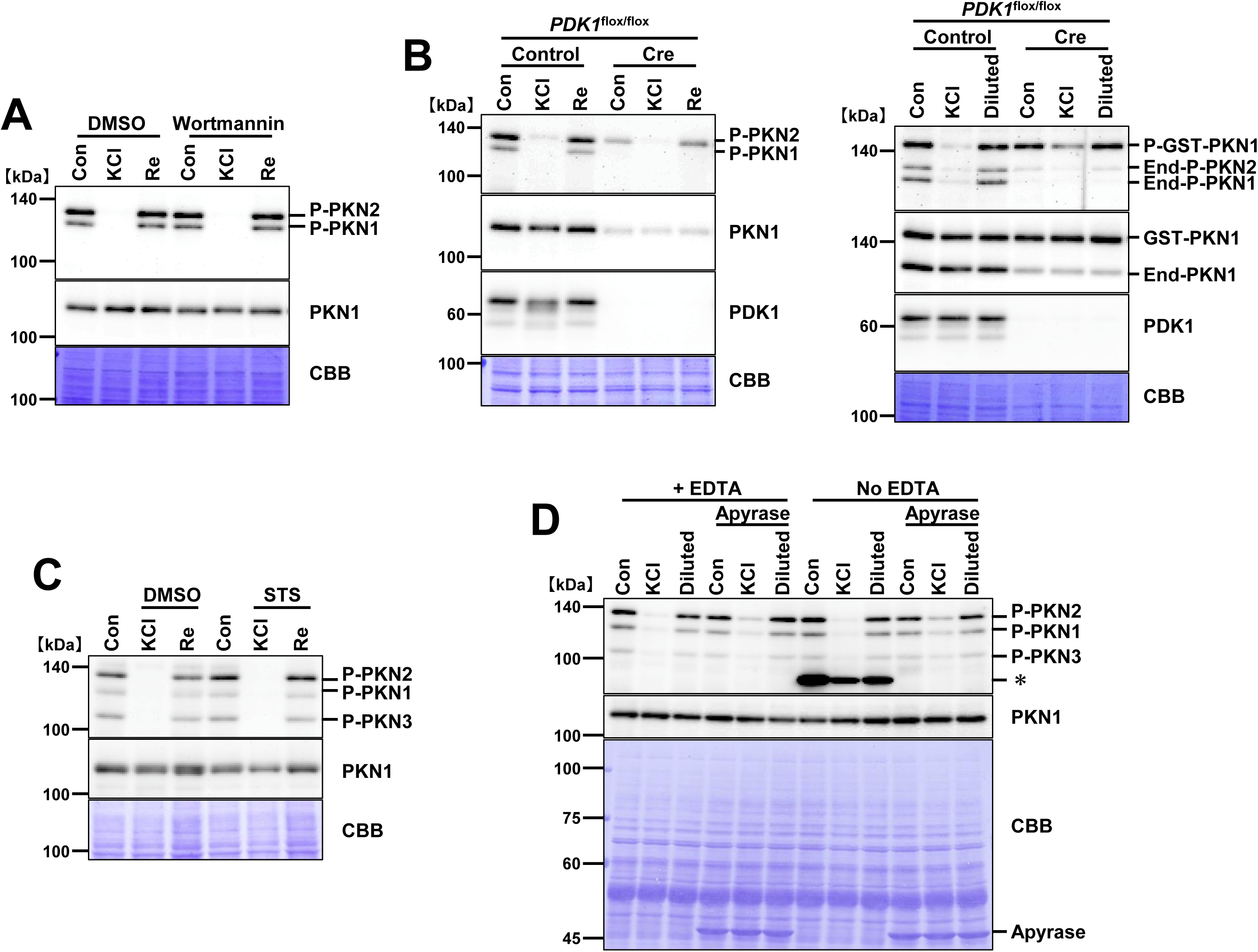
Recovery of PKN activation-loop phosphorylation after normalization of K⁺ concentration does not require PDK1, ATP or Mg²⁺. Whole-cell lysates were analyzed by immunoblotting with antibodies to phospho-PKN (activation loop) and total PKN; membranes were stained with Coomassie Brilliant Blue (CBB) as a loading control. Representative immunoblots are shown. “Con,” “KCl,” and “Re” denote the control, high-K⁺ treatment, and recovery in cell-based experiments, achieved by replacing the high-K⁺ medium with the original medium containing normal K⁺. In in-lysate experiments, “Diluted” indicates a 1:1 dilution into lysis buffer to lower ionic strength. (A) PI3K–PDK1 pathway inhibition with wortmannin. *PDK1*^flox/flox^ MEFs were pre-incubated with DMSO or 2 μM wortmannin for 3 h at 37°C (5% CO₂), then treated with 150 mM KCl for 10 min, followed by 5 min in normal-K⁺ medium. Wortmannin (or DMSO) was present throughout both phases. (B) PDK1 ablation by Cre and in-lysate reconstitution. *PDK1*^flox/flox^ MEFs were infected with control adenovirus (PDK1 present) or adenovirus expressing Cre recombinase (PDK1 knockout). After 48 h, cells were treated with 150 mM KCl for 10 min, then incubated in normal-K⁺ medium for 5 min (left panel). For the in-lysate assays (right panel), GST-tagged full-length PKN1 purified from Sf9 cells was incubated with soluble extracts from control virus–infected MEFs (PDK1 present) or from Cre-infected MEFs (PDK1 knockout) in the presence of 150 mM KCl for 10 min, then diluted 1:1 with lysis buffer prior to analysis. “End” indicates endogenous; “Diluted” marks the dilution step. (C) Broad-spectrum kinase inhibition with staurosporine. COS7 cells were treated with 150 mM KCl for 10 min in the presence of DMSO or 1 μM staurosporine, followed by 5 min in normal-K⁺ medium. Staurosporine (or DMSO) was maintained throughout. “STS” indicates staurosporine. (D) ATP- and Mg²⁺ -independent recovery of PKN activation-loop phosphorylation. COS7 soluble extracts were pre-incubated with 1.5 μM apyrase (recombinant, *E. coli*) with or without 1 mM EDTA for 1 h at 30°C to deplete endogenous ATP, then exposed to 150 mM KCl for 10 min; apyrase (and EDTA, when present) was maintained during the KCl incubation. Samples were then diluted 1:1 with lysis buffer prior to immunoblotting for phospho- and total PKN; CBB staining served as a loading control. Asterisks (*) denote non-specific signals observed in extracts that lacked both EDTA and apyrase. The following figure supplement is available for figure 4: **Figure supplement 1.** Autophosphorylation of PKN1 and recovery of activation-loop phosphorylation.

Next, we generated PDK1-knockout MEFs (PDK1-KO) by infecting *PDK1*^flox/flox^ MEFs with an adenovirus encoding Cre recombinase, and used MEFs infected with a control vector as a reference. In PDK1-KO MEFs, the expression of PKN1 was markedly reduced (Figure 4B), consistent with previous reports suggesting that activation-loop phosphorylation contributes to the stability of PKN1 (Balendran et al., 2000; Collins et al., 2005). Due to the low expression of PKN1 in PDK1-KO MEFs, it was difficult to evaluate its phosphorylation dynamics. Nevertheless, K⁺-induced decrease and subsequent recovery of PKN2 activation-loop phosphorylation were still observed in the absence of PDK1 (Figure 4B). To overcome the limitation posed by the low abundance of PKN1, we employed a cell-free system using full-length GST-tagged PKN1 phosphorylated at the activation loop, which was purified from Sf9 cells. The phosphorylated PKN1 protein was incubated with soluble extracts from PDK1-KO MEFs. Following treatment with KCl, the mixture was diluted twofold to reduce the K⁺ concentration. This procedure led to a recovery of activation-loop phosphorylation in PKN1, further confirming that the recovery mechanism is independent of PDK1 (Figure 4B).

### Recovery of PKN activation-loop phosphorylation is not dependent on autophosphorylation activity

We next investigated whether PKN1 autophosphorylation activity contributes to the recovery of activation-loop phosphorylation. Full-length GST-PKN1 purified from Sf9 cells is known to possess kinase activity (Yoshinaga et al., 1999). We incubated this active GST-PKN1 with non-phosphorylated GST-PKN1 fragments (GST-PKN1 aa 543-942, corresponding to the full catalytic domain, and GST-PKN1 aa 760-873, containing the activation-loop region) purified from *Escherichia coli* (*E. coli*) in the presence of ATP and Mg²⁺ at 30°C for 1 hour. Immunoblot analysis revealed that Thr774 in the activation loop of the PKN1 fragments expressed in *E. coli* was phosphorylated by active GST-PKN1 purified from Sf9 cells (Figure 4 - figure supplement 1A), suggesting that active PKN1 has its autophosphorylation activity at Thr774 *in vitro*.

To further examine whether PKN1 autophosphorylation is required for the recovery of activation-loop phosphorylation, we prepared fibroblast cell lines from a PKN1^T778A^ knock-in mouse model, in which the activation-loop Thr is substituted with Ala, resulting in complete loss of PKN1 kinase activity (Mehruba et al., 2020). These cells were transfected with one of the following three expression vectors for: (i) wild-type PKN1, (ii) kinase-negative PKN1[K644E] (mutation in the ATP-binding site), or (iii) PKN1[S916A] (mutation in the turn motif phosphorylation site, critical for kinase activity)(Lim et al., 2005). Since PKN1^T778A^ cells lack endogenous kinase activity, the autophosphorylation activity of PKN1 is expected to be observed only in cells transfected with vector (i) which expresses the active form of PKN1. Following high KCl treatment and subsequent return to normal medium, phosphorylation recovery was observed in all (i) (ii) (iii) conditions (Figure 4 - figure supplement 1B), suggesting that PKN1 autophosphorylation activity is not required for this process.

Furthermore, we investigated the role of other PKN isoforms by using His-Iα, a protein known to inhibit all PKN family members (Shiga et al., 2009; Yoshinaga et al., 1999). His-Iα was added to the soluble cell extracts from *PDK1*^flox/flox^ MEFs before KCl treatment, and after the decrease in PKN activation-loop phosphorylation, the reaction mixture was diluted twofold. Phosphorylation levels recovered to a similar extent in the presence or absence of His-Iα (Figure 4 - figure supplement 1C). This indicates that the recovery of the phosphorylation level after a decrease in K⁺ concentration does not require the intrinsic kinase activity of PKN family members.

### Recovery of activation-loop phosphorylation occurs without ATP or Mg²⁺ and is unaffected by broad-spectrum kinase inhibitors

We next examined whether the recovery of phosphorylation levels was influenced by kinase inhibitors. COS7 cells were incubated in medium containing staurosporine, a broad-spectrum kinase inhibitor, along with KCl. After treatment, the cells were returned to normal medium containing staurosporine, and activation-loop phosphorylation was assessed. Even in the presence of up to 1 µM staurosporine, phosphorylation levels recovered as in control conditions (Figure 4C).

Additionally, to assess the dependency on divalent cations, we chelated Mg²⁺ and Mn²⁺ in soluble extracts of COS7 cells by adding excess EDTA. After KCl treatment, the reaction mixture was diluted twofold, and phosphorylation levels were examined. Notably, recovery of PKN1 activation-loop phosphorylation was observed even in the presence of EDTA, suggesting that this process occurs independently of Mg²⁺ and Mn²⁺(Figure 4D).

Finally, we examined whether ATP was required for the recovery of phosphorylation levels. Soluble extracts of COS7 cells were treated with apyrase—an ATP diphosphohydrolase that hydrolyzes ATP to ADP and AMP—effectively depleting ATP from the reaction mixture (Hamasaki et al., 2009). These extracts were then incubated with 150 mM KCl, followed by a twofold dilution to lower the KCl concentration. As a result, activation-loop phosphorylation was recovered in ATP-depleted extracts to a comparable level as in ATP-containing controls (Figure 4D).

These findings demonstrate that the recovery of PKN activation-loop phosphorylation following a decrease in K⁺ concentration does not require PDK1, ATP, Mg²⁺, or intrinsic kinase activity of PKN. Instead, an alternative phosphotransfer mechanism, distinct from conventional kinase-mediated phosphorylation reactions, appears to regulate this process.

### A 14-residue peptide fragment encompassing the PKN1 activation-loop phosphorylation site (human PKN1 aa 767-780) exhibits phosphorylation loss upon high K⁺ treatment, followed by recovery upon reduction in K⁺ concentration

To identify the essential region within PKN1 responsible for the K⁺-induced decrease and subsequent recovery of activation-loop phosphorylation, we generated various deletion mutants of the PKN1 catalytic domain (Figure 5A). These constructs were expressed in COS7 cells, which were then treated with high KCl concentrations followed by incubation in normal medium. Immunoblot analysis revealed that the decrease and subsequent recovery of PKN1 activation-loop phosphorylation were observed in the PKN1 aa 689-942 fragment, indicating that the N-terminal region upstream of 688 aa is not required for this process (Figure 5B). To further narrow down the critical region, we generated shorter deletion mutants fused to GST and co-expressed them with PDK1 in *E. coli*. These fusion proteins were purified using GSH-Sepharose, allowing us to obtain pre-phosphorylated deletion mutant fragments. These phosphorylated fragments were then incubated with soluble extracts of COS7 cells and subjected to high KCl treatment. Immunoblot analysis showed that the GST-PKN1 aa 767-780 fragment exhibited a decrease in phosphorylation levels upon exposure to K⁺, which was reversed by twofold dilution of the reaction mixture to lower the K⁺ concentration (Figure 5C). To exclude the possibility that the observed phenomena were due to the use of PDK1 in generating the phosphorylated fragment, we employed the method described by Zhang et al., which uses genetic code expansion with orthogonal aminoacyl-tRNA synthetase–tRNA pairs to incorporate phosphothreonine directly into proteins without relying on kinases such as PDK1 (Zhang et al., 2017). In addition, to rule out any potential artifacts caused by the GST tag, we used a version of the activation-loop fragment bearing a (His)₆ tag for affinity purification. Consistent with previous results, this fragment also showed K⁺-induced decrease in phosphorylation followed by recovery upon K⁺ reduction (Figure 5 - figure supplement 1).

**Figure 5.**
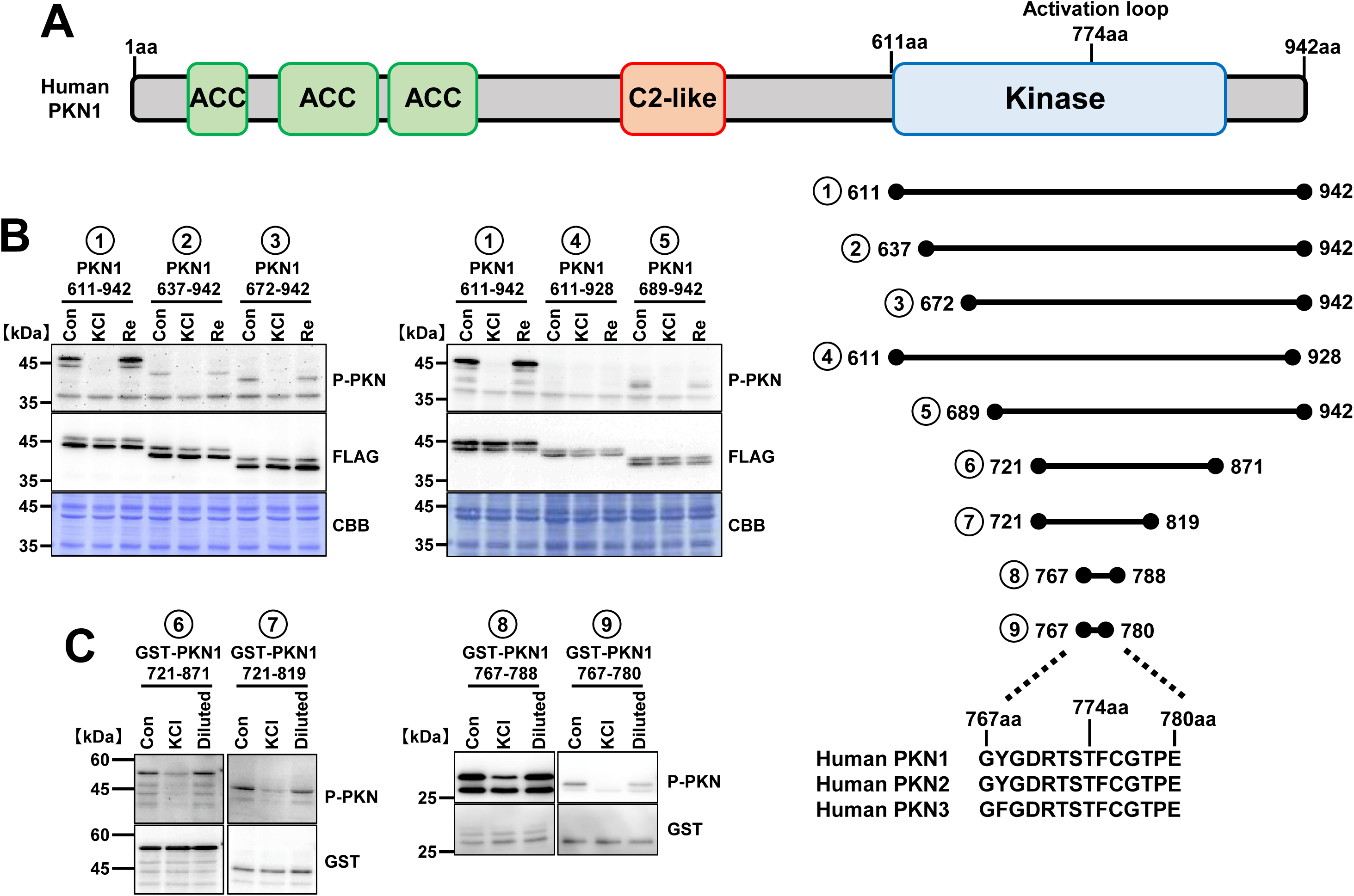
KCl-induced modulation of activation-loop phosphorylation is preserved in a minimal PKN1 fragment (aa 767–780). “Re” denotes the recovery phase in cell-based assays, achieved by replacing the high-K⁺ medium with the original medium containing normal K⁺. In lysate assays, “Diluted” indicates a 1:1 dilution into lysis buffer to reduce ionic strength. Blots were probed with antibodies to phospho-PKN (activation loop) and FLAG or GST where indicated; membranes were stained with CBB as a loading control. Representative immunoblots are shown. (A) Schematic of PKN1 and deletion constructs. Domain organization of human PKN1 with the activation loop indicated; the aa 767– 780 segment and Thr774 are marked. Sequence alignment around this region for PKN1/2/3 is shown (conserved Thr in the activation loop). (B) PKN1 deletion mutants expressed in COS7 cells. COS7 cells were transfected with plasmids encoding FLAG-tagged PKN1 mutants (aa 611–942, 637–942, 672–942, 611–928, and 689–942). After 48 h, cells were treated with 150 mM KCl for 10 min, then switched to normal-K⁺ medium for 5 min (“Re”). (C) Minimal fragments retain KCl responsiveness in lysates. GST-PKN1 721–871 and 721–819 were phosphorylated *in vitro* with PDK1 (3 h). GST-PKN1 767–788 and 767–780 were prepared by co-expression with PDK1. Phosphorylated fragments were mixed with COS7 cell extracts and incubated with or without 150 mM KCl for 10 min; “Diluted” samples were obtained by 1:1 mixing with lysis buffer before analysis. The following figure supplement is available for figure 5: **Figure supplement 1.** Effect of affinity tag on the recovery of PKN1 activation-loop phosphorylation.

Collectively, these findings demonstrate that the K⁺-induced decrease and subsequent recovery of activation-loop phosphorylation can occur within a minimal 14-residue region of PKN1 (aa 767–780) that includes the essential phosphothreonine site.

### The PKN1 activation loop loses its phosphate group upon high K⁺ concentration but reacquires the same phosphate group upon reduction in K⁺ concentration

After treatment with high K⁺ concentrations, a subsequent decrease in extracellular K⁺ levels rapidly recovered the activation-loop phosphorylation. Three mechanistic models could explain this recovery (Figure 6A):

**Figure 6.**
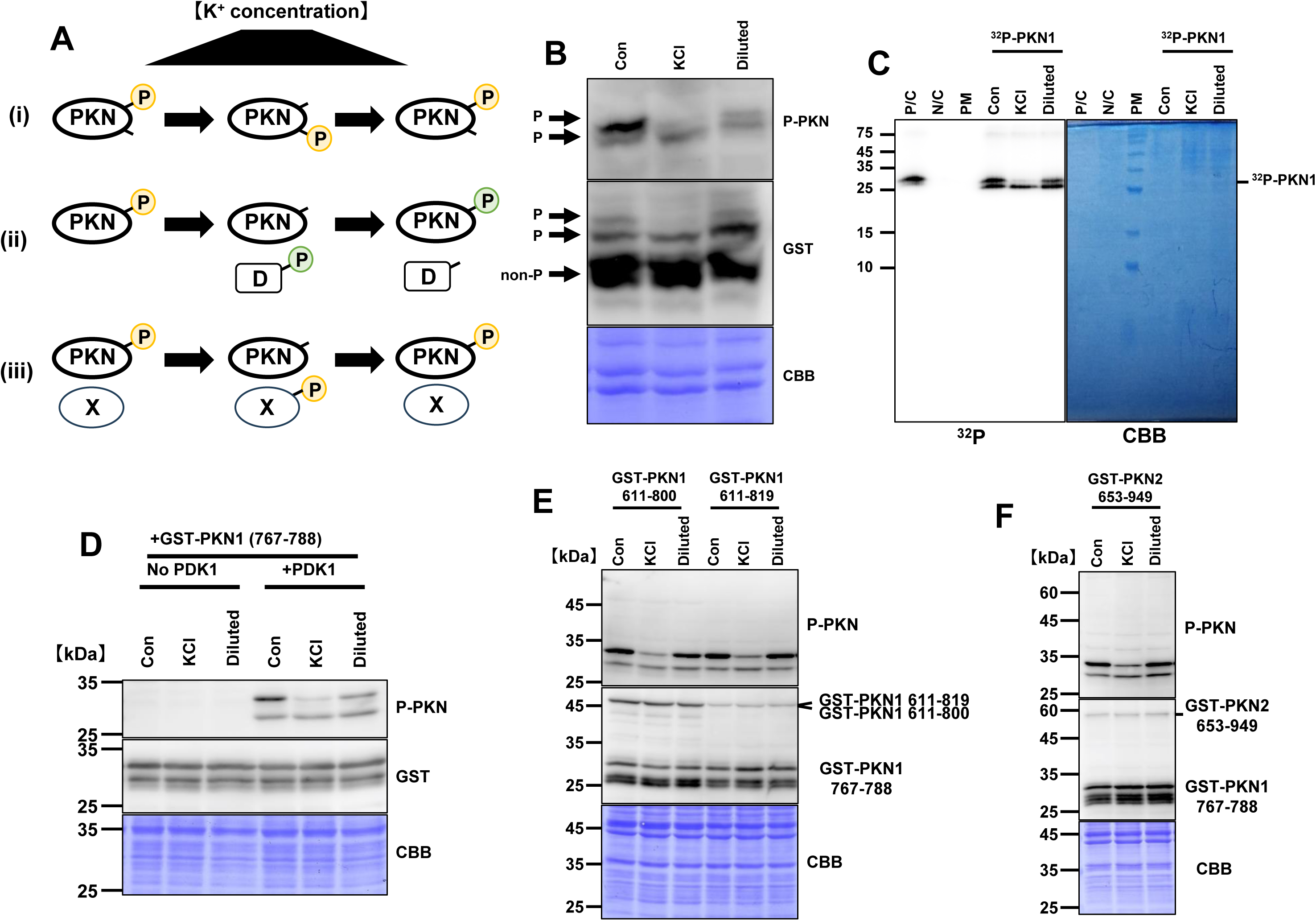
The phosphate initially used to modify the PKN1 activation loop is reused (recycled) during recovery. Unless otherwise noted, “Con,” “KCl,” and “Diluted” indicate the control condition, incubation with 150 mM KCl for 10 min, and subsequent 1:1 dilution into 1× lysis buffer to lower ionic strength, respectively. Immunoblots were probed with antibodies against phospho-PKN (activation loop) and GST where indicated; membranes were stained with CBB as a loading control. Representative immunoblots/autoradiographs are shown where applicable. (A) Working models for recovery. Three mechanistic models are considered: (i) intramolecular phosphate transfer within PKN; (ii) transfer from an external phosphate donor (factor D); and (iii) transient transfer to an intermediate (factor X) followed by return to PKN. Yellow “P” indicates the phosphate initially attached to the PKN activation loop; green “P” denotes a phosphate derived from factor D. (B) Detection of phosphorylated PKN1 using Phos-tag SDS-PAGE. COS7 cell extracts were incubated with phosphorylated GST–PKN1 (aa 767–788) in the presence of 150 mM KCl for 10 min; “Diluted” samples were generated by 1:1 mixing with 1× lysis buffer prior to analysis by Phos-tag SDS–PAGE and immunoblotting for phospho-PKN and GST. Bands labeled “P / non-P” indicate phosphorylated/non-phosphorylated species. (C) Detection of radiolabeled phospho-PKN1 by autoradiography. COS7 cell extracts were incubated with ^32^P-labeled phospho–GST–PKN1 (aa 767–788) under the same KCl and dilution conditions as in (B), followed by autoradiography to detect ^32^P– PKN1. P/C (positive control): radiolabeled phospho–GST–PKN1 alone; N/C (negative control): COS7 extract alone; PM: protein marker. (D) Recovery requires pre-existing activation-loop phosphorylation. GST–PKN1 (aa 767–788) purified from *E. coli* was preincubated with or without PDK1, then exposed to COS7 soluble extracts + 150 mM KCl for 10 min, followed by 1:1 dilution as above. Samples were immunoblotted for phospho-PKN and GST. (Only PDK1-pretreated fragments support recovery.) (E) Selectivity of PKN1 fragments undergoing recovery of activation-loop phosphorylation. COS7 extracts containing phospho–GST–PKN1 (aa 767–788) were mixed with non-phosphorylated GST–PKN1 (aa 611–819) or GST–PKN1 (aa 611–800) and processed as in (B). (F) Isoform selectivity. COS7 extracts containing phospho–GST–PKN1 (aa 767–788) were incubated with non-phosphorylated GST–PKN2 (aa 653–949) and analyzed as in (B). The following figure supplements are available for figure 6: **Figure supplement 1.** Preserved K⁺-dependent loss–recovery dynamics in activation-loop region point mutants. **Figure supplement 2.** ATP-independent recovery of PKN1 activation-loop phosphorylation. **Figure supplement 3.** Absence of recovery with dephosphorylated PKN1 fragment.

#### Model (i): Intramolecular transfer of phosphate group

The phosphate group on the PKN1 activation loop is transferred to another amino acid residue within the same molecule upon high K⁺ treatment. When K⁺ levels decrease, the phosphate group is transferred back to the original activation-loop residue (Figure 6A, i).

#### Model (ii): Transfer of phosphate group from an external donor

Upon reduction in K⁺concentration, the activation loop of PKN1 is phosphorylated by an unidentified phosphate group donor (D), distinct from ATP or PKN1 itself (Figure 6A, ii).

#### Model (iii): Reversible transfer of phosphate group

High K⁺ induces the transfer of the phosphate group from the PKN1 activation loop to an intermediary factor (X). When K⁺ levels decrease, the phosphate group is transferred back to the activation loop (Figure 6A, iii). To distinguish among these models, we analyzed the phosphorylated status of the PKN1 activation loop using Phos-tag assays. Phos-tag binds phosphate groups and alters the electrophoretic mobility of phosphorylated proteins in SDS-PAGE (Kinoshita et al., 2022; Kinoshita et al., 2006). When phosphorylated PKN1 aa 767-788 was incubated with soluble extract of COS7 cells and subjected to high KCl treatment, the Phos-tag signal corresponding to phosphorylated PKN1 decreased (Figure 6B). Following dilution of the reaction mixture to reduce K⁺ levels, the phosphorylated PKN1 band was recovered. These results indicate that the number of phosphate groups attached to the PKN1 activation loop decreases upon high K⁺ treatment, ruling out the model (i).

One scenario consistent with model (i) is that, under high K⁺, the phosphate group is transiently relocated to another amino-acid residue but is then lost during SDS–PAGE sample preparation and electrophoresis, so that the transferred phosphate cannot be detected. Based on the experiment in Figure 5C using the PKN1 767–780 pT774 peptide, any such acceptor would have to reside within the activation loop itself. Notably, the PKN1 activation loop sequence (GYGDRTSTFCGTPE) contains a cysteine at P+2 (C776). Cysteine phosphorylation in the activation segment of PINK1 kinase has recently been reported (Waddell et al., 2023), and pCys is known to be heat- and acid-labile, raising the possibility that a pCys intermediate could be hydrolyzed in samples subjected to standard SDS–PAGE handling. To test this, we generated a C776A mutant of PKN1 and examined the response to high-K⁺ treatment followed by ion dilution. As shown in Figure 6 - figure supplement 1A, the basal anti-pT774 signal in C776A was weaker than in wild-type; nevertheless, high K⁺ still induced a decrease in activation-loop threonine phosphorylation, and subsequent K⁺ reduction recovered the phosphorylation, mirroring the wild-type behavior. We performed analogous point-mutation experiments in Akt1, which also harbors a cysteine at P+2 (C310) in its activation-loop (Figure 6 - figure supplement 1B), and found that the C310A mutant likewise displayed the same high-K⁺–induced decrease and post-dilution recovery (Figure 6 - figure supplement 1A). We obtained indistinguishable results whether samples were boiled or not prior to SDS–PAGE, arguing against artifactual loss of phosphate during sample preparation (data not shown). Given that noncanonical, acid-labile side-chain phosphorylations could in principle occur—e.g., N-phospho-arginine (phosphoramidate) and acyl-phosphate on aspartate (and, theoretically, glutamate)—we substituted Asp770, Arg771, and Glu780 in the PKN1 activation loop (Thr774 activation segment) with alanine (D770A, R771A, E780A), noting that E780 lies within the conserved APE motif and substitutions here may also perturb activation-segment architecture. Upon high-K⁺ treatment and subsequent dilution, all three mutants exhibited the same pattern of activation loop phospho-Thr decrease and recovery as the wild-type (Figure 6 - figure supplement 1C). Collectively, these results argue against model (i), in which the pT774 phosphate is reversibly transferred to another residue within the activation loop (e.g., C776) and subsequently lost due to heat/acid lability during SDS–PAGE experiments. We further examined the fate of the phosphate group by conducting radioactive phosphate group tracing experiments. Non-phosphorylated GST-PKN1 aa 767-788, purified from *E. coli*, was incubated with [γ-³²P]ATP, Mg²⁺, and recombinant PDK1 to radioactively label its activation-loop phosphorylation. After free ATP was removed using GSH-Sepharose purification, the phosphorylated PKN1 fragment was incubated with soluble extract of COS7 cells. The mixture was subjected to high KCl treatment, followed by dilution to reduce K⁺ levels. Autoradiography revealed that the radioactive signal on the PKN1 band decreased upon high K⁺ treatment and was recovered after K⁺ concentration was reduced (Figure 6C). This finding suggests that the phosphate group transiently dissociates from the PKN1 activation loop when K⁺ levels are high and is reattached when K⁺ concentration is lowered, further ruling out the model (i). If model (ii) were valid, this would imply the presence of a radiolabeled phosphate group donor (factor D). However, since [γ-³²P]ATP was not added to the K⁺ treatment reaction mixture containing mammalian soluble cell extract, the presence of a radiolabeled phosphorylated factor D is unlikely. Nevertheless, we cannot entirely exclude the possibility that residual [γ-³²P]ATP carried over into the KCl-treated reaction mixture interacted with factor D present in the cell extract, leading to the generation of a radiolabeled factor D within the reaction system. To eliminate this possibility, we performed the same experiment in the presence of excess cold ATP, which would compete with any residual radioactive ATP. Even under these conditions, autoradiography showed that the loss and subsequent recovery of phosphorylation occurred as before (Figure 6 - figure supplement 2), making the model (ii) unlikely.

Taken together, the results from Phos-tag assays and radioactive phosphate group tracing experiments support the model (iii), in which an intermediary factor (X) transiently accepts the phosphate group from the PKN1 activation loop when K⁺ concentrations are high and returns it upon reduction in K⁺ concentration.

### Phosphate group dynamics require pre-existing phosphorylation of the PKN1 activation loop

If model (iii) is correct, then PKN1 fragments that were never phosphorylated should be unable to undergo phosphate group transfer upon K⁺ treatment. To test this hypothesis, we prepared non-phosphorylated PKN1 activation-loop fragments from *E. coli* and compared them with PDK1-phosphorylated fragments. Each fragment was incubated with soluble extract of COS7 cells and subjected to high K⁺ treatment followed by dilution. Immunoblot analysis revealed that only the phosphorylated PKN1 fragment experienced dissociation of phosphate groups and subsequent reassociation, whereas the non-phosphorylated fragment remained unmodified throughout (Figure 6D). Similarly, when phosphorylated GST-PKN1 aa 767-788 was treated with λ-phosphatase *in vitro* to remove its phosphate group, subsequent incubation in soluble extract of COS7 cells followed by high K⁺ treatment and dilution failed to recover phosphate group attachment (Figure 6 - figure supplement 3). These results are consistent with the prediction of the model (iii).

### Phosphate group transfer does not occur between distinct PKN molecules

If the intermediary factor X was a freely diffusing phosphate group carrier, it should be capable of transferring phosphate groups between different PKN1 molecules. To test this, we co-incubated phosphorylated GST-PKN1 aa 767-788 with non-phosphorylated GST-PKN1 aa 611-800 or GST-PKN1 aa 611-819 (expressed in *E. coli* without PDK1 phosphorylation) in soluble cell extract of COS7 cells, followed by high K⁺ treatment and dilution. Immunoblot analysis showed that only the originally phosphorylated GST-PKN1 aa 767-788 underwent reversible phosphorylation loss, while non-phosphorylated GST-PKN1 aa 611-800 and GST-PKN1 aa 611-819 did not acquire phosphate groups (Figure 6E). Furthermore, when phosphorylated GST-PKN1 aa 767-788 was co-incubated with non-phosphorylated PKN2 aa 653-949, no phosphate group transfer was observed between them following K⁺ treatment and dilution (Figure 6F). These results suggest that the phosphate group transfer mechanism involves a specific interaction between the activation loop of PKN1 and the intermediary factor X, rather than a general phosphate group exchange among PKN family members.

Our findings demonstrate that the PKN1 activation loop transiently loses its phosphate group upon high K⁺ exposure but reacquires the same phosphate group when K⁺ levels return to normal. This process does not involve conventional kinase-mediated phosphorylation or a free phosphate group donor but instead relies on an intermediary factor (X) that transiently accepts and subsequently returns the phosphate group. These results reveal a novel phosphorylation homeostasis mechanism that is distinct from traditional kinase-phosphatase regulation.

## Discussion

### Implications of the observations at both the cellular and cell-free levels

In this study, we demonstrated that changes in extracellular Na⁺ and K⁺ concentrations, as well as alterations in these ion concentrations in cell-free extracts, induce changes in the activation-loop phosphorylation in multiple AGC kinases. Although the precise molecular mechanisms underlying these phenomena remain unclear, several lines of evidence suggest that a common mechanism is likely involved:

i. Previous studies have reported that increasing extracellular Na⁺ and K⁺ concentrations to levels used in our experiments also leads to an increase in intracellular ion concentrations (Pleinis et al., 2021; Walz & Hertz, 1983; Wojnowski & Oberleithner, 1991).
ii. In both cell-based and cell-free systems, an increase in ion concentration led to a decrease in activation-loop phosphorylation of PKN and other kinases, and this decrease was suppressed by the application of PP2A inhibitors.
iii. In both cell-based and cell-free systems, phosphorylation levels recovered after reducing the ion concentration following high-ion treatment, and this recovery was not suppressed even in the presence of broad-spectrum kinase inhibitors.

These findings suggest the existence of a pathway in which Na⁺ and K⁺ regulate the activity of intracellular protein kinases independently of changes in cell volume or membrane potential. Furthermore, this phenomenon was observed across all tested eukaryotic lines, including human embryonic kidney HEK293 cells, human cervical cancer HeLa cells, human neuroblastoma SH-SY5Y cells, African green monkey kidney COS7 cells, mouse embryonic fibroblasts (MEFs), and insect Sf9 cells. However, this response was not observed in *E. coli* cell extracts (Figure 1 - figure supplement 2), suggesting that this mechanism may be an evolutionarily conserved feature in eukaryotic cells. Notably, K⁺ modulated activation-loop phosphorylation even at concentrations relatively close to its physiological intracellular levels. Eil et al. reported that tumor interstitial fluid contains K⁺ concentrations exceeding 40 mM and that increasing extracellular K⁺ levels to 40 mM elevates intracellular K⁺ concentration to 153 mM (Eil et al., 2016). These findings suggest that localized increases in K⁺ concentrations in pathological conditions such as tumors or tissue damage may be sufficient to induce changes in kinase phosphorylation levels. Given that activation-loop phosphorylation is essential for kinase activity, this regulatory mechanism likely has significant implications for various cellular functions.

### Known intracellular targets of Na⁺ and K⁺ in signal transduction and their possible relevance to activation-loop dynamics

To date, there are only a few signaling molecules that have been identified as intracellular targets of Na⁺ and K⁺. One of the well-characterized examples is the WNK (With No Lysine) kinase family, which is known to be inhibited by Cl⁻ ions. Interestingly, WNK kinases can also bind K⁺ ions in a Cl⁻-independent manner, and direct inhibition of WNK activity by K⁺ has been suggested (Goldsmith & Rodan, 2023; Pleinis et al., 2021; Rodan, 2023). WNK4 activity is most strongly inhibited at K⁺ concentrations ranging from 80 to 180 mM. Furthermore, increasing the K⁺ concentration in the culture medium (more than 50 mM) has been reported to rapidly elevate intracellular K⁺ levels, leading to the suppression of intracellular WNK activity. However, unlike the changes of AGC kinase activation-loop phosphorylation reported in this study, WNK kinase activity is entirely ATP-dependent. This fundamental difference suggests that WNK kinases are unlikely to be directly involved in the observed Na⁺ or K⁺-induced regulation of AGC kinase activation-loop phosphorylation.

### Possible mechanisms of loss and reacquisition of phosphate groups

The activation-loop phosphorylation of AGC protein kinases has been widely accepted to be catalyzed by the master kinase PDK1. However, in this study, we demonstrated that the reacquisition of phosphate group in the PKN activation loop, which occurs upon a decrease in K⁺ concentration following an initial increase, proceeds independently of PDK1, ATP, and Mg²⁺. In the phosphotransfer model proposed in model (iii) of Figure 6A, an unidentified factor X is hypothesized to acquire phosphate group from the PKN activation loop, functioning as a phosphate group donor independent of ATP. Potential candidates for phosphorylated factor X include alternative phosphate group donors present in mammalian soluble cell extracts, such as guanosine triphosphate (GTP), uridine triphosphate (UTP), cytidine triphosphate (CTP), phosphoenolpyruvate (PEP), phosphocreatine (PCr), inorganic polyphosphate (polyP), inositol heptakisphosphate (IP7)/inositol octakisphosphate (IP8), and 1,3-bisphosphoglycerate (1,3-BPG). Some of these molecules, including GTP (Bostrom et al., 2009), PEP (Dong et al., 2016), IP7/IP8 (Saiardi et al., 2004), and 1,3-BPG (Morino et al., 1991; Oslund et al., 2017), have been reported to donate phosphate groups to proteins either enzymatically or non-enzymatically. However, none of these phosphate group donors are known to directly acquire phosphate groups from phosphorylated proteins. These candidates require ATP-dependent phosphorylation for their generation except for 1,3-BPG. Given the absence of [γ-³²P]ATP and the presence of excess cold ATP in the reaction mixture (Figure 6C), it is unlikely that radiolabeled phosphate group donors were synthesized in our study. Therefore, these candidate molecules are unlikely to serve as phosphorylated factor X. 1,3-BPG is unique in that it can be synthesized from inorganic phosphate (Pi) and glyceraldehyde-3-phosphate (G3P) by glyceraldehyde-3-phosphate dehydrogenase (GAPDH), even in the absence of ATP (Zhang et al., 2021). Thus, one possible explanation is that radiolabeled Pi released from phosphorylated activation-loop fragments by some phosphatase such as PP2A is incorporated into 1,3-BPG in the reaction mixture, which subsequently functions as phosphorylated factor X. However, mammalian soluble cell extracts contain high levels of endogenous “cold” Pi stably (typically 1–10 mM (Bevington et al., 1986) and the reaction mixture in our study contained 0.1 - 1 mM. Given this, it is unlikely that radiolabeled Pi (estimated to be at most 1 μM in the reaction mixture) is selectively and efficiently incorporated into 1,3-BPG and subsequently utilized for activation-loop phosphorylation.

An alternative possibility is that factor X is a protein and that phosphate group transfer occurs between proteins. If this is the case, initially radiolabeling the phosphate group on the PKN activation loop should allow visualization of phosphorylated factor X via SDS-PAGE upon K⁺ treatment. However, autoradiography following 20% SDS-PAGE did not reveal any distinct bands corresponding to phosphorylated factor X (Figure 6C). One possible explanation is that factor X is a low-molecular-weight peptide or a protein that cannot be efficiently resolved by gel electrophoresis. Our experimental system demonstrated that, following high K⁺ treatment and subsequent reduction in K⁺ concentration, only those activation-loop fragments that originally contained a phosphate group were capable of reacquiring it. This suggests that reacquisition of phosphate group is selective and may be mediated by a close interaction between the activation-loop fragment and factor X. A similar mechanism has been described in Saccharomyces cerevisiae, where the Sln1-Ypd1-Ssk1 pathway relays osmotic stress signals via sequential phosphotransfer. In this system, specific interactions between proteins ensure highly selective and efficient phosphotransfer, including bidirectional phosphate group exchange between the aspartate residue of Sln1 and the histidine residue of Ypd1 (Janiak-Spens et al., 2005). However, in our study, the phosphate groups involved in PKN activation-loop regulation are attached to serine or threonine residues. Compared to the phospho-histidine or phospho-aspartate linkages in bacterial and yeast phosphotransfers, phospho-serine and phospho-threonine bonds are chemically more stable. This suggests that transferring phosphate groups from serine or threonine to another molecule and then back to serine or threonine would be chemically challenging. Furthermore, given the universal reliance on ATP-dependent phosphorylation systems in mammalian cells, it raises the question of why an ATP-independent phosphate group transfer mechanism would be evolutionarily necessary. To date, no such mechanism has been reported in mammalian cells to our knowledge.

### Biological implications of an ATP-independent phosphate group dynamics

Nevertheless, the existence of an ATP-independent phosphotransfer system, as demonstrated in this study, suggests that precise control of activation-loop phosphorylation levels is critical for cell survival. Whether physiologically or pathologically induced, changes in Na⁺ or K⁺ concentrations significantly affect the activity of multiple kinases, which in turn influence cellular functions such as proliferation, motility, and cell cycle regulation. Given this, the presence of a robust and selective phosphate group recovery system is plausible. An ATP-independent phosphate group reacquisition mechanism would provide an energy-efficient means of regulating kinase activity, particularly under conditions of metabolic stress. Moreover, selectively recovering only those activation-loop phosphorylation that were originally lost, rather than inducing an overall increase in phosphorylation, could prevent excessive activation of multiple kinases and mitigate potential pathological consequences. This mechanism could represent an effective homeostatic strategy for maintaining kinase activity within an optimal range.

### Potential role of PP2A in phosphate group transfer

Factor X may not only function as a phosphate group donor/acceptor but also possess enzymatic activity for phosphotransfer or interact with a phosphotransfer enzyme. In the yeast Sln1-Ypd1-Ssk1 system mentioned earlier, Sln1 itself has phosphotransferase enzyme activity from Sln1 to Ypd1, while Ypd1 facilitates phosphotransfer from Ypd1 to Sln1 without intrinsic catalytic activity.

In our study, we demonstrated that PP2A is necessary for the phosphate group loss from the activation loop upon Na⁺ or K⁺ elevation. PP2A is widely recognized for its conventional protein phosphatase activity, catalyzing the hydrolysis of phosphoserine and phosphothreonine residues to release Pi. If a PP2A isoform possesses phosphotransfer enzymatic activity (which could be inhibited by rubratoxin and okadaic acid), then a simple model could be proposed where PP2A mediates phosphate group transfer between the activation loop and factor X. However, no prior studies have reported phosphotransfer activity for PP2A. Alternatively, PP2A may regulate an unidentified phosphotransfer enzyme by modulating its phosphorylation state. If such a phosphotransfer enzyme depends on phosphorylation for its activity, PP2A might control its function via dephosphorylation. However, this model seems to be inconsistent with our experimental observations, as kinase inhibitors or ATP depletion did not prevent phosphate group transfer in our system. Another possibility is that PP2A functions as a scaffold or adaptor that facilitates interactions between the activation loop and a phosphotransfer enzyme. PP2A has been reported to dephosphorylate the catalytic domain phosphorylation sites of various AGC kinases, thereby regulating their enzymatic activity, and some of these interactions have been demonstrated through co-immunoprecipitation experiments (Bielinski & Mumby, 2007; Hansra et al., 1996; Kuo et al., 2008; Liauw & Steinberg, 1996; Rodgers et al., 2011). If PP2A lacks intrinsic phosphotransfer activity but associates with a phosphotransfer enzyme, it could facilitate phosphate group transfer between the activation loop and factor X. Na⁺ and K⁺ concentrations may influence the association of these protein molecules or directly affect the activity of the phosphotransfer enzyme. At present, the molecular identity of factor X and the precise mechanism of the phosphotransfer remain unknown. However, our findings suggest that AGC kinase-mediated signaling may be subject to a novel mode of regulation involving distinct phosphotransfer mechanisms. Further investigations are required to elucidate these mechanisms and their potential physiological relevance.

## Acknowledgements

The authors thank Masumi Eto for the helpful discussion.

## Materials and Methods

### Cell culture

COS7 cells, HEK293 cells, HeLa cells, and *PDK1*^flox/flox^ MEFs were cultured in Dulbecco’s Modified Eagle Medium (DMEM; Nacalai Tesque, Kyoto, Japan) supplemented with 10% fetal bovine serum (FBS; Gibco), 100 μg/mL penicillin (Nacalai Tesque), and 100 U/mL streptomycin (Nacalai Tesque) at 37°C in a 5% CO₂ incubator. Primary *PDK1*^flox/flox^ mouse embryonic fibroblasts (MEFs) were kindly provided by Dr. Ogawa, Kobe University. Primary wild-type (WT) MEFs derived from 14.5-day-old embryos of wild-type mice were prepared as previously described (Danno et al., 2017). Adenoviruses expressing Cre recombinase (AxCANCre) were prepared as previously described (Danno et al., 2017). *PDK1*^flox/flox^ MEFs were infected with AxCANCre for 2 days to induce PDK1 deletion. PKN1[T778A] MEFs, described previously (Mashud et al., 2017; Mehruba et al., 2020), were immortalized by retroviral transduction of the SV40 large T antigen. These immortalized PKN1[T778A] MEFs were cultured in DMEM supplemented with 10% FBS and maintained at 37°C in a 5% CO₂ incubator. Sf9 cells were cultured in Sf-900™ III SFM (Gibco) supplemented with 5% FBS, 1% Antibiotic-Antimycotic (100X; Gibco), and 6.8 mM glutamic acid at 28°C in an incubator.

### Primer information for making PKN1 and Akt1 deletion mutant

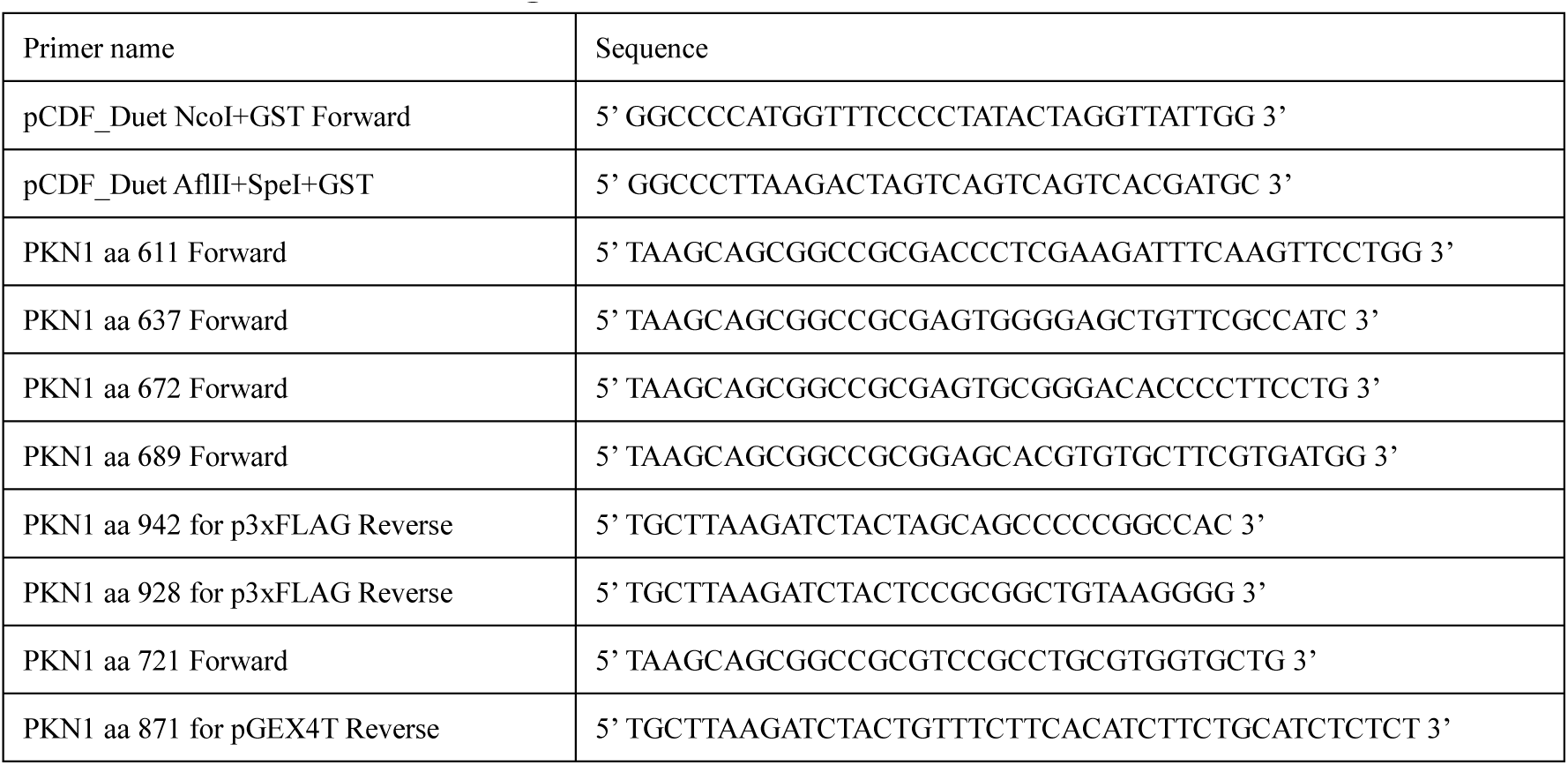
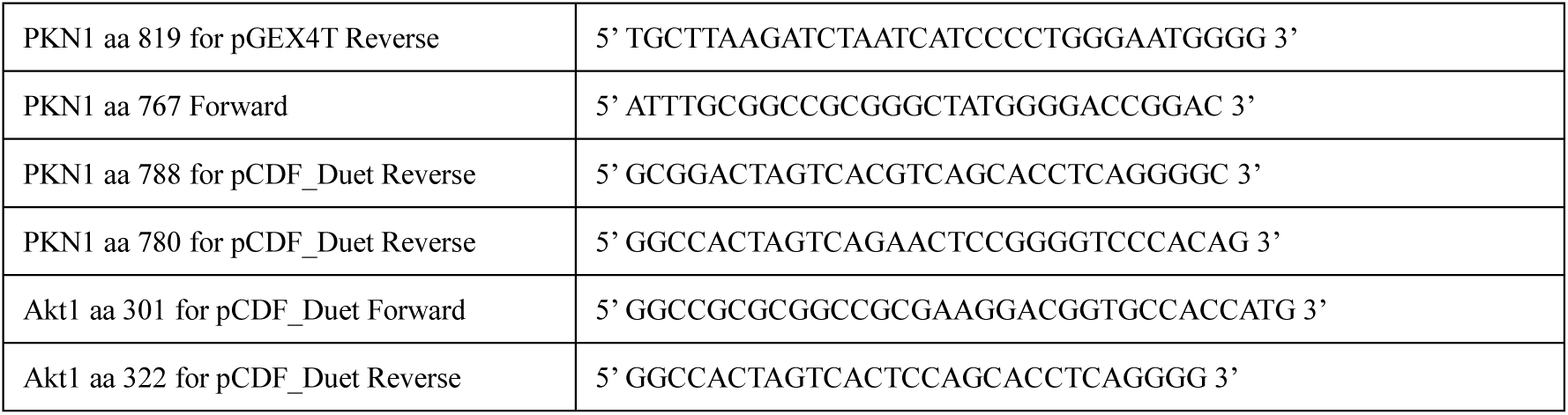

### Plasmid constructs

Vectors for expressing recombinant proteins in mammalian cells were constructed as follows: Insert fragments encoding PKN1 aa 611-942, aa 637-942, aa 672-942, aa 689-942, and aa 611-928 were amplified by PCR from human PKN1 DNA (Yoshinaga et al., 1999) and subcloned into the *Not*I/*Bgl*II site of the p3xFLAG CMV vector. Vectors for expressing recombinant proteins in *E. Coli* were constructed similarly. pGEX4T-HB vector was constructed using the pGEX4T vector as a backbone, in which its multiple cloning site (MCS) was replaced with the MCS from the p3xFLAG-CMV-10 vector. Insert fragment encoding PKN1 aa 721-871 and aa 721-819 for the expression of PKN1 deletion mutants were amplified by PCR from human PKN1 DNA, and subcloned into the *Not*I/*Bgl*II site of pGEX4T-HB vector. Insert DNA *Nco*I-GST-*Not*I-*Afl*II was amplified by PCR from pGEX-6p-2 and subcloned into the *Nco*I/*Afl*II site of the pCDF_Duet containing PDK1 gene kindly provided by Dr. O’Donoghue, University of Western Ontario, London (Balasuriya et al., 2018). Insert fragments encoding PKN1 aa 767-788 and aa 767-780 were amplified and subcloned into the *Not*I/*Spe*I site of the pCDF_Duet PDK1 co-expression vector, which containing the *Nco*I-GST-*Not*I-*Spe*I. Alanine-substitution point mutants were generated by site-directed mutagenesis—PKN1: G769A, D770A, R771A, C776A, T778A, E780A; Akt1: C310A, T312A—using the PrimeSTAR Mutagenesis Basal Kit with PrimeSTAR Max DNA Polymerase (Takara Bio) according to the manufacturer’s instructions. All constructs were verified by Sanger sequencing.

### Expression of PKN1 deletion mutants in COS7 cells

COS7 cells were cultured in antibiotic-free DMEM supplemented with 10% FBS and transfected with 3 μg plasmid DNA per 6-cm dish using HilyMax (Dojindo). After 3 h, the medium was replaced with antibiotic-free DMEM supplemented with 10% FBS. Cells were assayed 48 h post-transfection (for example, treated with 150 mM KCl for 10 min and then incubated in normal-K^+^ medium for 5 min for recovery).

### Expression and purification of recombinant PKNs in insect cells and *E. coli*

Protein expression and purification from Sf9 cells and *E. coli* were performed as previously described (Takahashi et al., 1999; Yoshinaga et al., 1999). To express PKN1 aa 689–800 phosphorylated at Thr774 within the activation loop in *E. coli* using genetic code expansion technique without co-expression of PDK1, we used BL21(DE3) ΔserC ΔycdX transformed with the plasmid pUC_pThrRS_SeptRNAv2.0CUA_OXB20-PduX_EFSep, as described in (Zhang et al., 2017). The plasmid was kindly provided by Dr. Jason W Chin, MRC Laboratory of Molecular Biology. Protein expression and purification were performed according to the methods described in (Zhang et al., 2017).

### RNA interference

Knockdown of PP2A catalytic subunit expression was performed using the following siRNAs: s10959 for human PPP2CA, s10961 for human PPP2CB, and Silencer® Select Negative Control #1 (Applied Biosystems/Ambion) as a negative control. HeLa cells cultured in 6-well plates were transfected with 10 nM siRNA using Lipofectamine RNAiMAX (Invitrogen).

### Treatment of cells or cell-derived preparations with ions

Cells were counted using a hemocytometer, and 1 × 10⁵ cells were plated into 6-cm dishes. After 24 hours, the culture medium was replaced with antibiotic-free DMEM containing either NaCl or KCl, and cells were incubated for 10 min at 37°C in a 5% CO_2_ incubator. Subsequently, the medium was replaced with antibiotic-free DMEM, and cells were incubated for 5 min under the same conditions. For NaCl- or KCl-treated samples at each step, the medium was removed, and the cells were immediately frozen in liquid nitrogen to quickly stop enzymatic reactions. The frozen cells were added with 1 x SDS-sample buffer and subjected to immunoblotting. For preparation of whole-cell lysates, cells were lysed in nine volumes of 1× lysis buffer containing 50 mM Tris–HCl (pH 7.5), 1 mM EGTA, 0.5 mM DTT, and 0.1% Triton X-100, then sonicated with a Branson Sonifier 250 (output level 4) for 20 s. Soluble extracts were obtained by centrifugation of whole-cell lysates at 20,400 x g for 30 min at 4°C. Whole-cell lysates or clarified soluble extracts were aliquoted into parallel sets as follows: two sets underwent NaCl/KCl treatment and one set served as a no-salt control. The two treated sets were incubated with NaCl or KCl (at the indicated concentrations) for 10 min at 30 °C. Thereafter, one treated set remained at high salt until quench (high-salt condition), whereas the other was diluted 1:1 with 1× lysis buffer and incubated for an additional 5 min at 30 °C before quench (post-dilution condition). No-salt controls were processed in parallel with 1× lysis buffer substituted for salt at each step. Volumes were matched at each step to keep the starting lysate/extract concentration identical across conditions, and all samples were finally brought to 1× SDS by adding an equal volume of 2× SDS sample buffer.

### Antibodies

Anti-phospho-S6K (T229) antibody was purchased from Abcam. Anti-PKCλ antibody (610207) and Anti-PDK1 antibody (611070) were purchased from BD Biosciences. Anti-phospho-PRK1 (Thr774)/PRK2 (Thr816) antibody (#2611), anti-p38 antibody (#9212), anti-phospho-p38 (Thr180/Tyr182) antibody (#4511), anti-phospho-PKCζ/λ(Thr410/403) antibody (#9378), anti-p70 S6 Kinase antibody (#9202), anti-PP2A C subunit antibody (#2259), anti-GAPDH (14C10) Rabbit mAb (#2118), anti-phospho-Akt (Thr308) Antibody (#9275) were purchased from Cell Signaling Technology. Anti-FLAG M2 antibody was purchased from Sigma. Anti-GST antibody (B-14) was purchased from SANTA CRUZ BIOTECHNOLOGY. The polyclonal antibody αC6 against PKN1 were prepared as previously described (Mukai et al., 1994). The phospho-specific antibodies used recognize activation-loop phosphosites: PKN1 Thr774 / PKN2 Thr816, PKCζ Thr410 / PKCλ Thr403, and p70 S6K Thr229 (as specified by the manufacturers’ datasheets and prior reports). This study’s “phospho-PKN/PKCζ/λ/S6K” signals therefore refer to activation-loop phosphorylation.

### Immunoblot analysis and Phos-tag SDS-PAGE

Cells or cell preparations with SDS-sample buffer were subjected to 8–20% SDS-PAGE. The separated proteins were transferred onto a polyvinylidene difluoride membrane. The membrane was blocked for 1 hour at room temperature using TBS-T (20 mM Tris-HCl, pH 7.5, 137 mM NaCl, and 0.05% Triton X-100) supplemented with 5% normal goat serum or Blocking One (Nacalai Tesque). Following blocking, the membrane was incubated with the primary antibody diluted in TBS-T for 1 hour at room temperature or overnight at 4°C. After primary antibody incubation, the membrane was washed three times with TBS-T (5 min per wash) and then incubated for 1 hour in TBS-T containing the secondary antibody conjugated to horseradish peroxidase at a dilution of 1:2,000 to 1:10,000. The membrane was subsequently washed three times with TBS-T (10 min per wash). Signal detection was performed using an enhanced chemiluminescence method.

Gels for Phos-tag SDS-PAGE were prepared with a stacking gel consisting of 4% (w/v) acrylamide, 125 mM Tris-HCl (pH 6.8), 0.1% (w/v) SDS, 0.2% (v/v) N,N,N′,N′-tetramethylethylenediamine (TEMED), and 0.08% (w/v) ammonium persulfate (APS). The separating gel was composed of 12% (w/v) acrylamide, 375 mM Tris-HCl (pH 8.8), 0.1% (w/v) SDS, 25 μM Phos-tag acrylamide, 50 μM MnCl₂, 0.1% (v/v) TEMED, and 0.05% (w/v) APS.

### *In vitro* phosphatase assay

Phospho-casein was prepared as a substrate for PP2A as previously described (Tallant & Cheung, 1984). The PP2A complex purified from Sf9 cells was incubated for 6 min at 30°C with phospho-casein in a reaction mixture containing 20 mM Tris-HCl (pH 7.5), 150 mM KCl or 600 mM NaCl. Inorganic phosphate release was measured as described (H. Mukai et al., 1993).

### Autoradiography analysis

GST-PKN1 aa 767-788 was incubated for 3 hours at 30°C in a reaction mixture containing 10 mM Tris-HCl (pH 7.5), 5 mM MgCl₂, 185 KBq of [γ-32P] ATP, and GST-PDK1 purified from *E. coli.* Decrease in radiolabeled GST-PKN1 aa 767-788 phosphorylation in COS7 cell lysate was induced by addition of 150 mM KCl, and recovery of the phosphorylation was triggered by a twofold (1:1) dilution. The reaction was terminated by adding SDS sample buffer, followed by boiling for 5 min. The reaction mixture was subjected to SDS-PAGE. The gel was fixed, dried, and analyzed via autoradiography using a Typhoon FLA 9500 imager (GE Healthcare Life Sciences).

### Preparation and assay of apyrase

The expression plasmid for apyrase was kindly provided by Dr. Kato (Jichi Medical University) (Hamasaki et al., 2009). *E. coli* BL21(DE3) cells were transformed with the plasmid and cultured in Luria-Bertani (LB) medium containing 50 μg/ml ampicillin. Recombinant apyrase expression was induced by adding isopropyl β-D-thiogalactoside (IPTG) to a final concentration of 200 μM, followed by overnight incubation at 28°C. The bacterial cells were resuspended in sonication buffer H (20 mM HEPES, pH 8.0, 100 mM NaCl, 8 M urea) and lysed by sonication. The lysate was centrifuged at 100,000 x g for 30 min, and the supernatant was incubated with Ni-NTA agarose beads for 1 hour at 4°C. The beads were washed three times with sonication buffer H supplemented with 20 mM imidazole. The lysate-resin mixture was loaded into an empty column with the bottom cap attached, and recombinant apyrase was eluted using sonication buffer H containing 300 mM imidazole. The eluted sample was loaded onto a PD-10 column pre-equilibrated with 20 mM HEPES (pH 7.4), and urea and imidazole were removed by gel filtration. Soluble apyrase was subsequently collected.

The phosphohydrolase activity of the purified apyrase was assayed by incubating the enzyme for 20 min at 37°C in a reaction mixture containing 10 mM Tris-HCl (pH 7.5), 9.25 kBq of [γ-³²P]ATP, and 1 mM cold ATP. The release of inorganic phosphate was quantified as described previously (H Mukai et al., 1993). The specific activity of the purified apyrase was approximately 3 mmol/min per g of protein.

### Statistical analysis

All experiments were independently performed at least three times. Statistical significance between two groups was determined using Student’s t-test, with a p-value < 0.01 considered significant. Data presented in the figures and text represent the mean ± standard error of the mean (SEM) of representative experiments unless otherwise stated.

## Data availability

The datasets generated and/or analyzed during the current study are available from the corresponding author upon reasonable request.

---

**Figure 1 – figure supplement 1.**
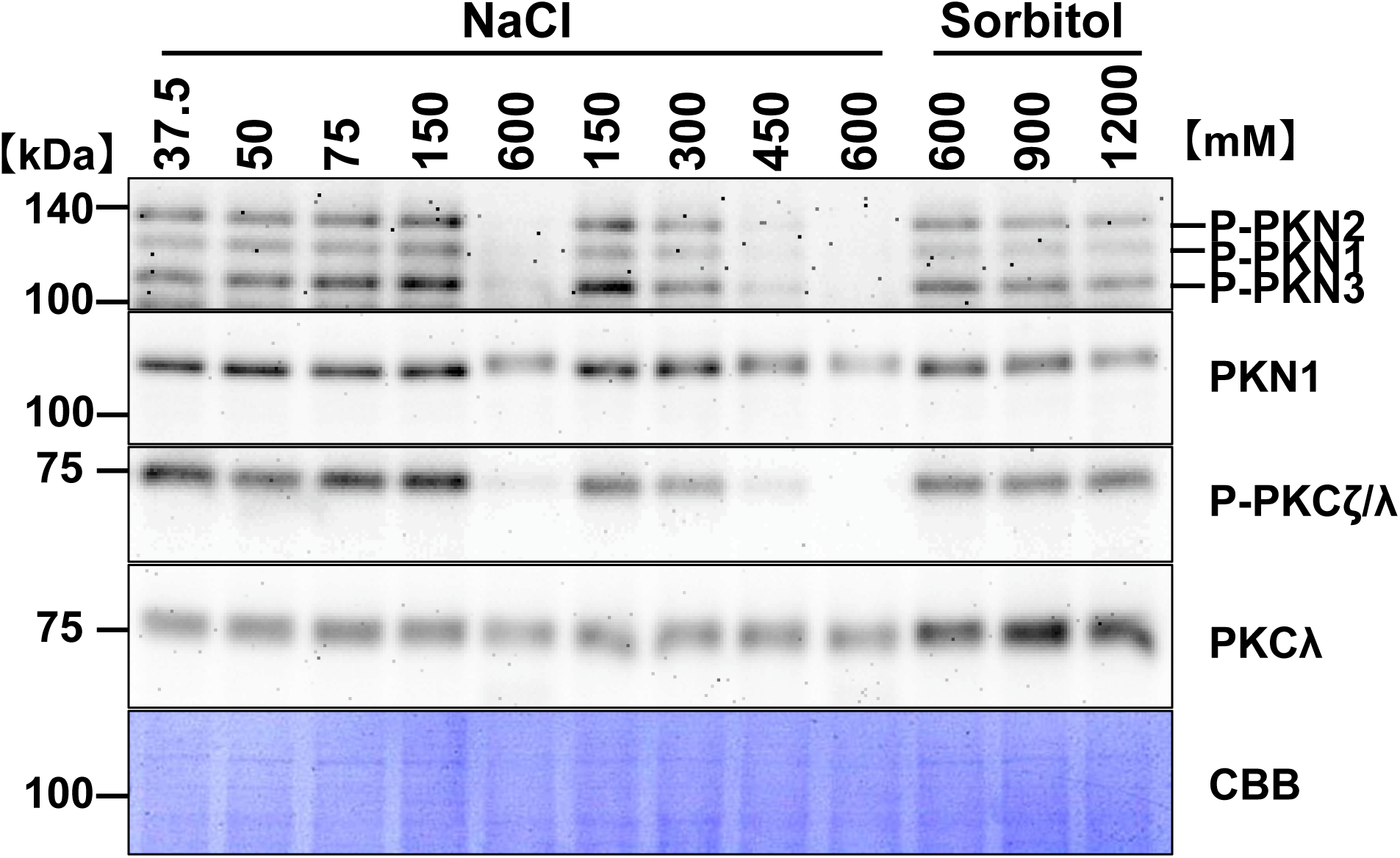
Effect of varying NaCl concentrations on activation-loop phosphorylation of PKN and PKCζ/λ in COS7 cells. COS7 cells were incubated for 10 min in a HEPES-based medium (4 mM KCl, 2.5 mM CaCl₂, 1 mM MgCl₂, 10 mM HEPES) supplemented with NaCl or sorbitol at the indicated concentrations (mM). Cell extracts were immunoblotted with antibodies against phospho-PKN (activation loop), total PKN, phospho-PKCζ/λ, and total PKCλ; membranes were stained with Coomassie Brilliant Blue (CBB) as a loading control. Representative immunoblots are shown.

**Figure 1 – figure supplement 2.**
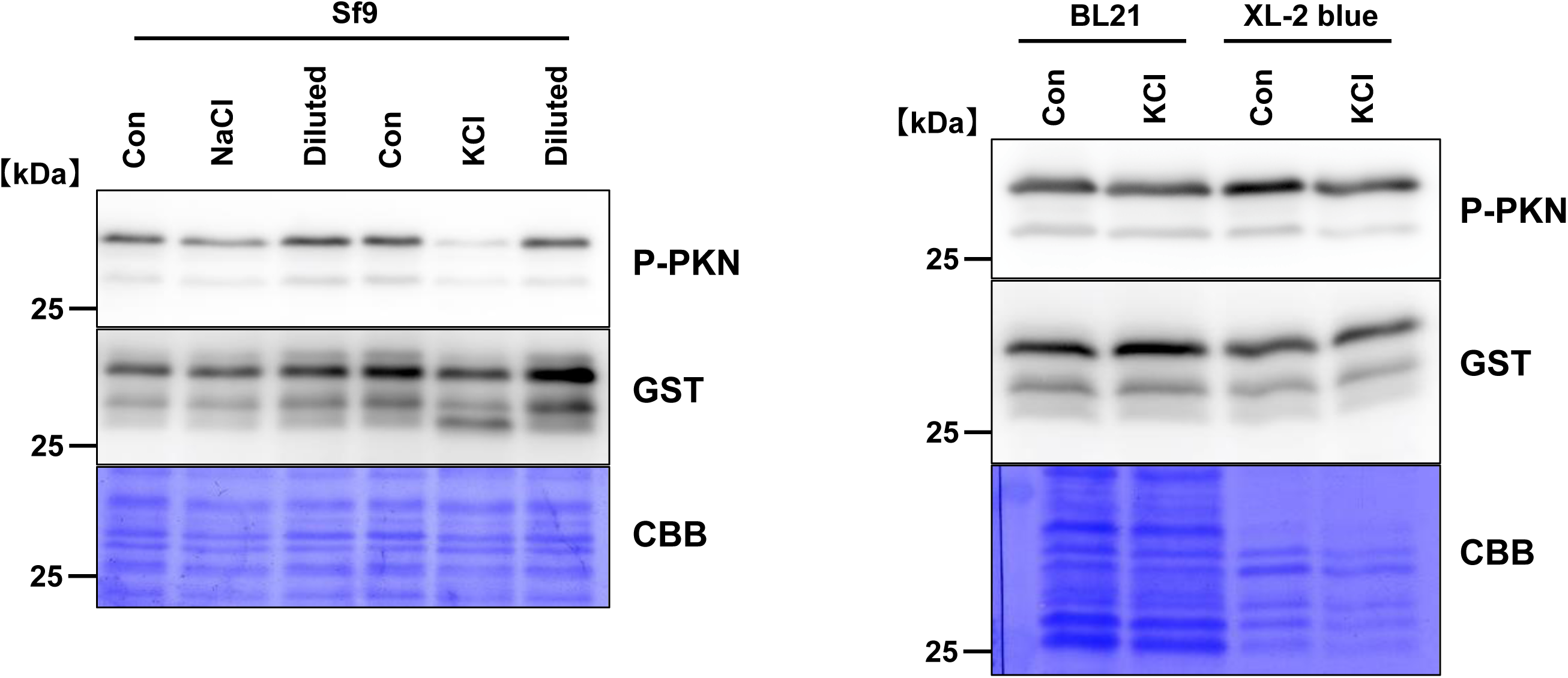
KCl-induced decrease in PKN phosphorylation in insect-cell extracts (Sf9) but not in *E. coli* extracts. Extracts from Sf9 cells and from *E. coli* strains (BL21 and XL1-Blue) containing phospho-GST–PKN1 (aa 767–788) were incubated with 600 mM NaCl or 150 mM KCl for 10 min. “Diluted” samples were prepared by 1:1 mixing with 1× lysis buffer (50 mM Tris-HCl pH 7.5, 1 mM EGTA, 0.5 mM DTT, 0.1% Triton X-100). Immunoblots were probed for phospho-PKN and GST. Mmembranes were stained with Coomassie Brilliant Blue (CBB) after immunoblotting as a loading control. Representative immunoblots are shown.

**Figure 2 – figure supplement 1.**
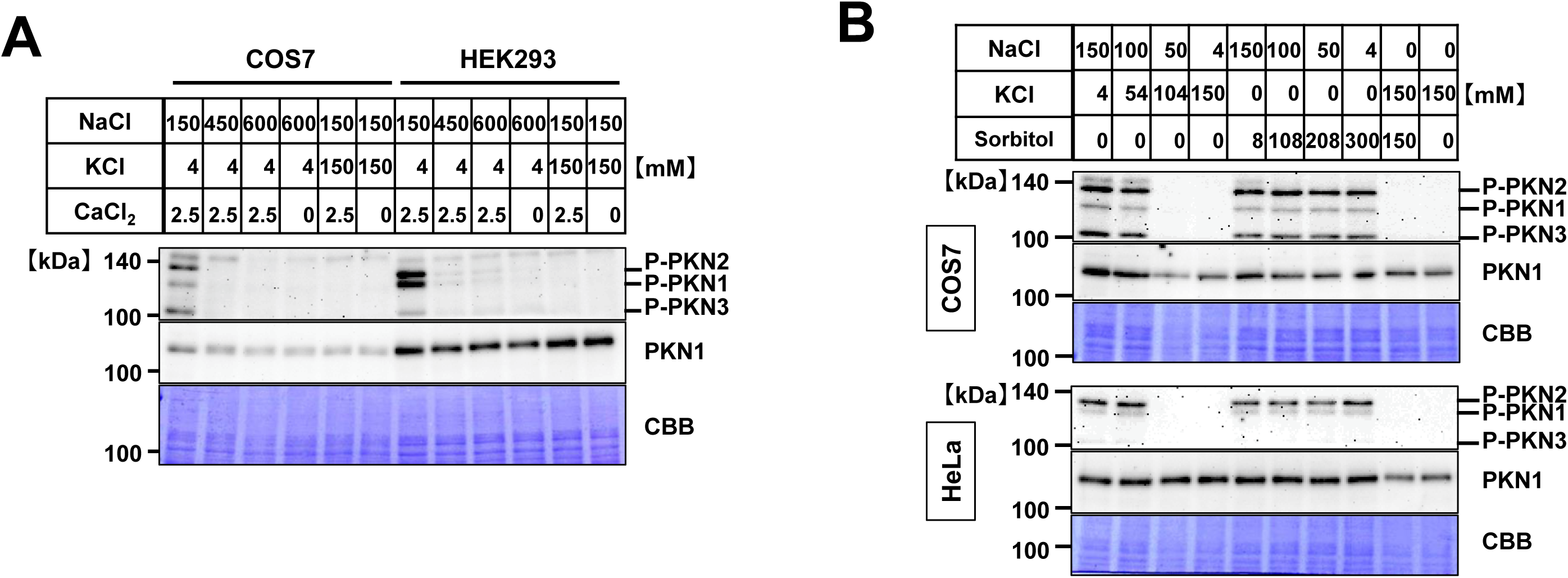
Effects of membrane potential and osmotic pressure on PKN activation-loop phosphorylation. Cell extracts were immunoblotted for phospho-PKN (activation loop) and total PKN; Coomassie Brilliant Blue (CBB) staining of the membrane after immunoblotting was used as a loading control. Representative blots are shown. (A) Ca²⁺ independence. COS7 and HEK293 cells were incubated for 10 min in a HEPES-based medium (10 mM HEPES, 1 mM MgCl₂) supplemented with NaCl, KCl, and CaCl₂ at the indicated mM. (B) Osmolality independence. COS7 and HeLa cells were incubated for 10 min in a HEPES-based medium (10 mM HEPES, 2.5 mM CaCl₂, 1 mM MgCl₂) supplemented with NaCl, KCl, or sorbitol at the indicated mM.

**Figure 4 – figure supplement 1.**
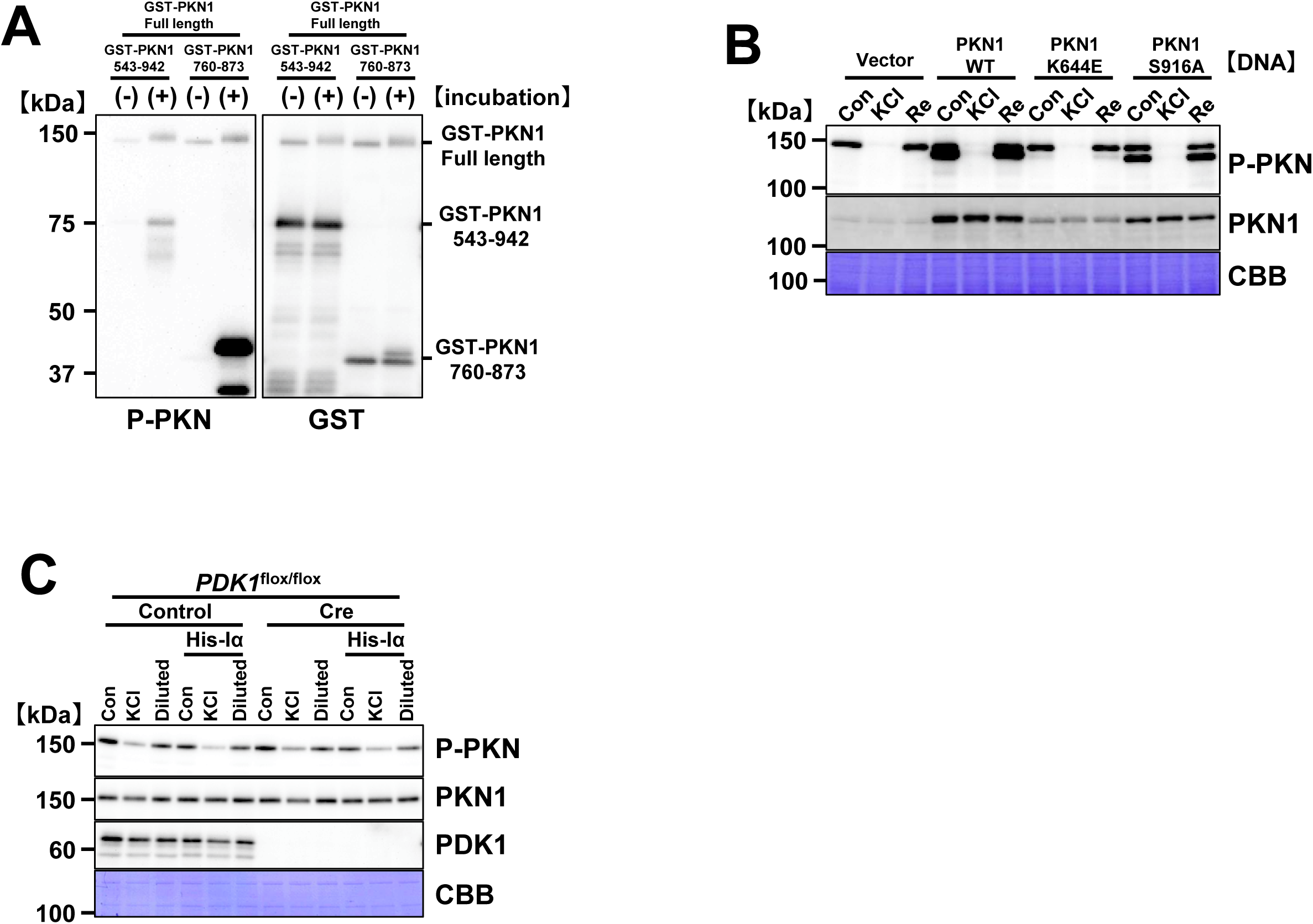
Autophosphorylation of PKN1 and recovery of activation-loop phosphorylation. Immunoblot analysis was performed using antibodies against the total and phosphorylated forms of PKN, GST and PDK1. Coomassie Brilliant Blue (CBB) staining of the membrane after immunoblotting was used as a loading control. (A) Autophosphorylation *in vitro*. Full-length GST–PKN1 purified from Sf9 and non-phosphorylated GST–PKN1 (aa 543–942, aa 760–873) purified from *E. coli* were incubated in 10 mM Tris-HCl (pH 7.5), 5 mM MgCl₂, 30 μM ATP at 30°C or 90 min. (B) Recovery independent of PKN1 catalytic activity in cells. Immortalized PKN1[T778A] MEFs were transfected with empty vector or PKN1 constructs (WT, K644E, S916A), treated with 150 mM KCl for 10 min, then switched to normal-K⁺ medium for 5 min to induce recovery. (C) His-Iα does not block recovery in lysates. Soluble extracts from *PDK1*^flox/flox^ MEFs ± PDK1 were incubated with full-length GST–PKN1 and His-Iα in 150 mM KCl for 10 min. “Diluted” samples were prepared by 1:1 mixing with 1× lysis buffer (50 mM Tris-HCl pH 7.5, 1 mM EGTA, 0.5 mM DTT, 0.1% Triton X-100).),

**Figure 5 – figure supplement 1.**
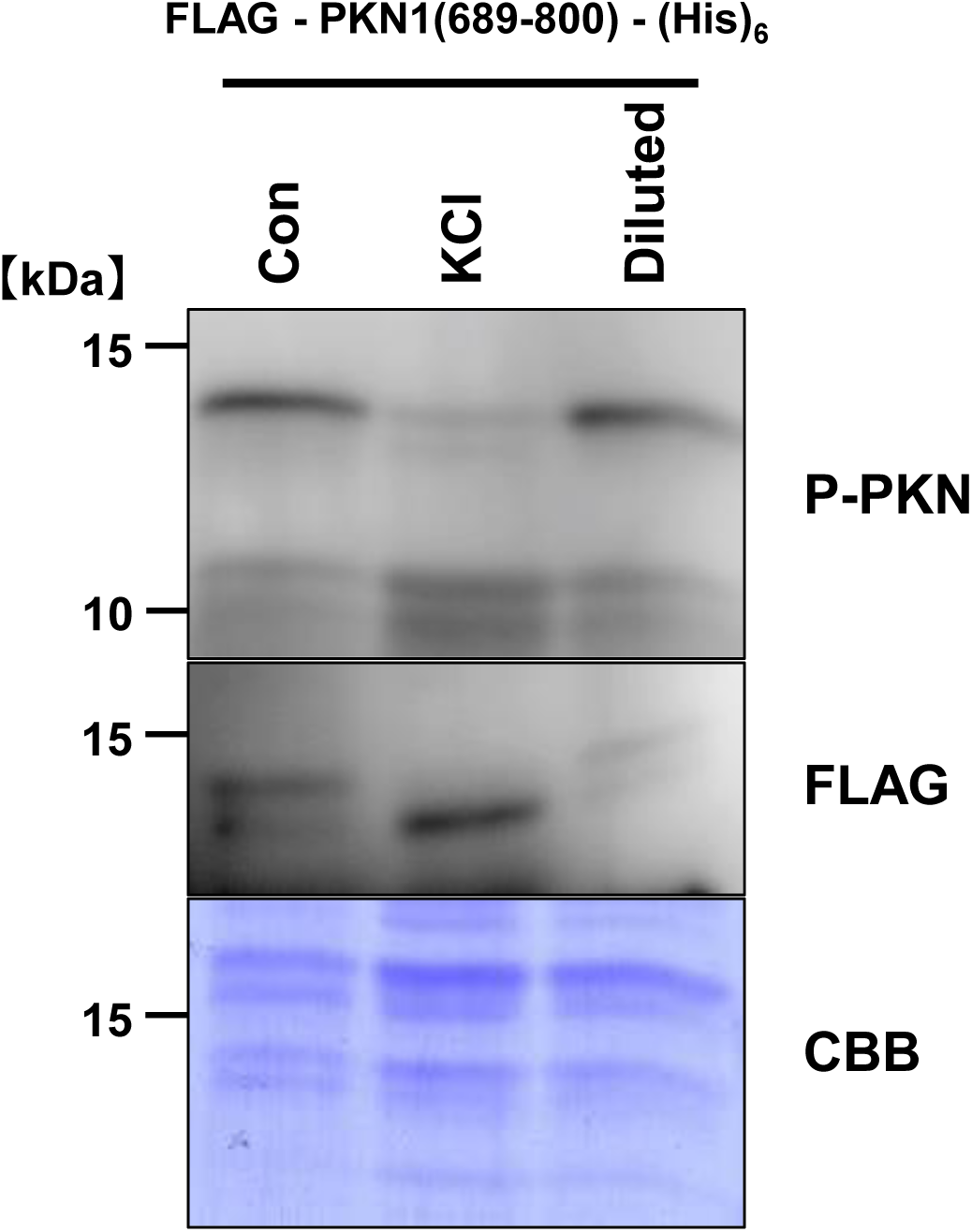
Effect of affinity tag on the recovery of PKN1 activation-loop phosphorylation. COS7 cell extracts were incubated with KCl and the FLAG and (His)_6_ - tagged proteins; “Diluted” samples were prepared by 1:1 mixing with 1× lysis buffer (50 mM Tris-HCl pH 7.5, 1 mM EGTA, 0.5 mM DTT, 0.1% Triton X-100). Immunoblots were probed with anti–phospho-PKN (activation loop) and anti-FLAG. Membranes were stained with Coomassie Brilliant Blue (CBB) after immunoblotting as a loading control.

**Figure 6 – figure supplement 1.**
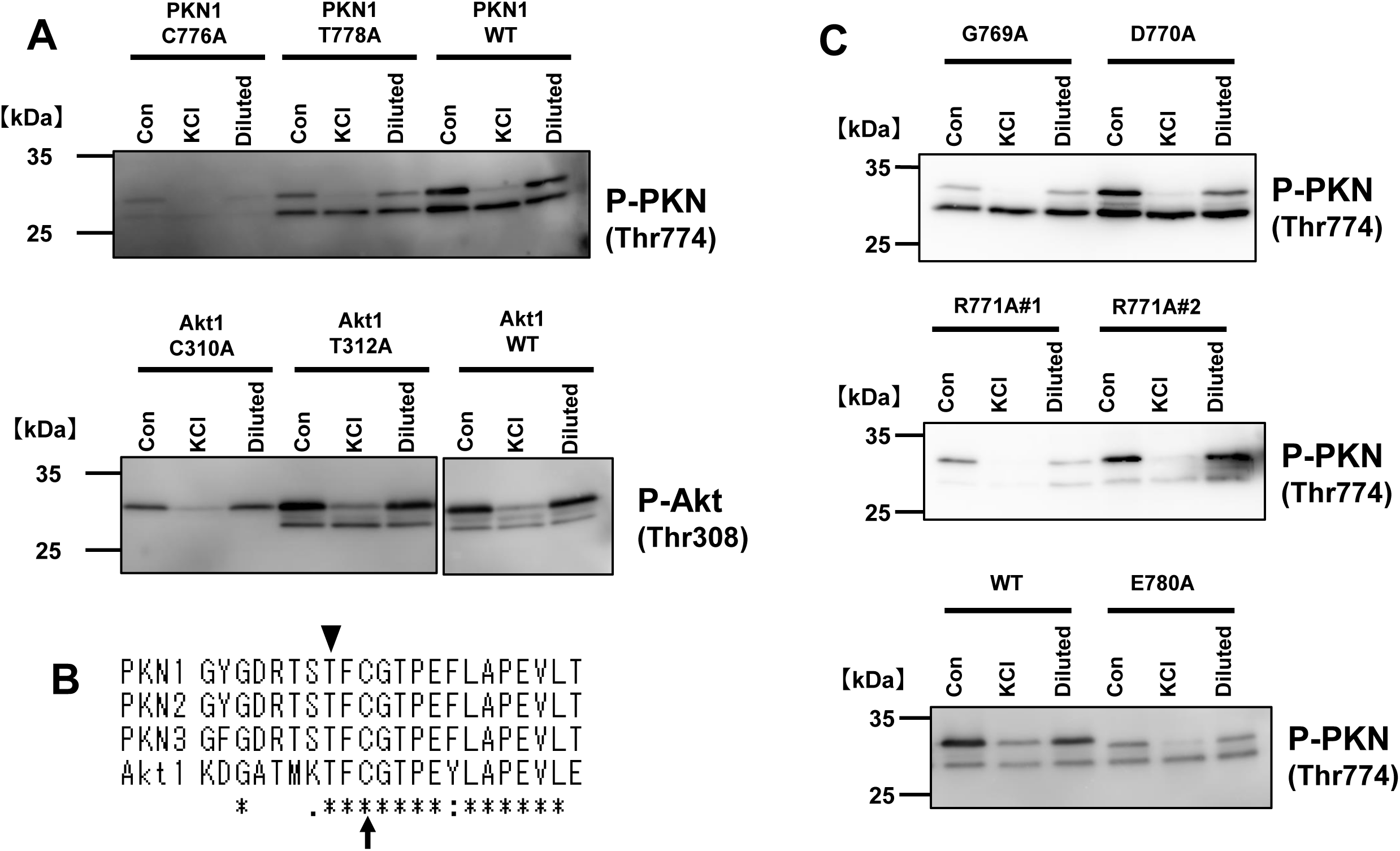
Preserved K⁺-dependent loss–recovery dynamics in activation-loop region point mutants. (A) Representative immunoblots showing activation-loop phosphorylation of PKN1 (P-PKN) and Akt1 (P-Akt) in wild-type (WT) and point-mutant constructs under three conditions: Con (control), KCl (high-K⁺ treatment), and Diluted (post-dilution, i.e., ion reduction). For GST-PKN1 (aa 767–788), mutants include C776A (the conserved Cys at P+2 within the TFCGT motif) and T778A; for GST-Akt1 (aa 301–322), mutants include C310A (the corresponding conserved Cys) and T312A. In all cases, high K⁺ reduces the activation-loop phosphor-signal, which recovers upon ion reduction, similarly to WT. Molecular weight markers (kDa) are indicated. (B) Alignment of activation-loop sequences from PKN1, PKN2, PKN3, and Akt1. The arrow indicates the conserved cysteine (Cys776 in PKN1; Cys310 in Akt1), and the arrowhead marks the activation-loop threonine that, when phosphorylated, is detected by the phospho-specific antibodies used in (A) and (C). (C) Additional GST-PKN1 (aa 767–788) point mutants at activation-loop residues with side chains that could, in principle, form labile phosphate linkages were tested: D770A, R771A (two clones), E780A, alongside WT. As in (A), high K⁺ reduces the activation-loop phospho-signal, which returns after dilution, indicating that these substitutions do not prevent the loss-and-reacquisition behavior.

**Figure 6 – figure supplement 2.**
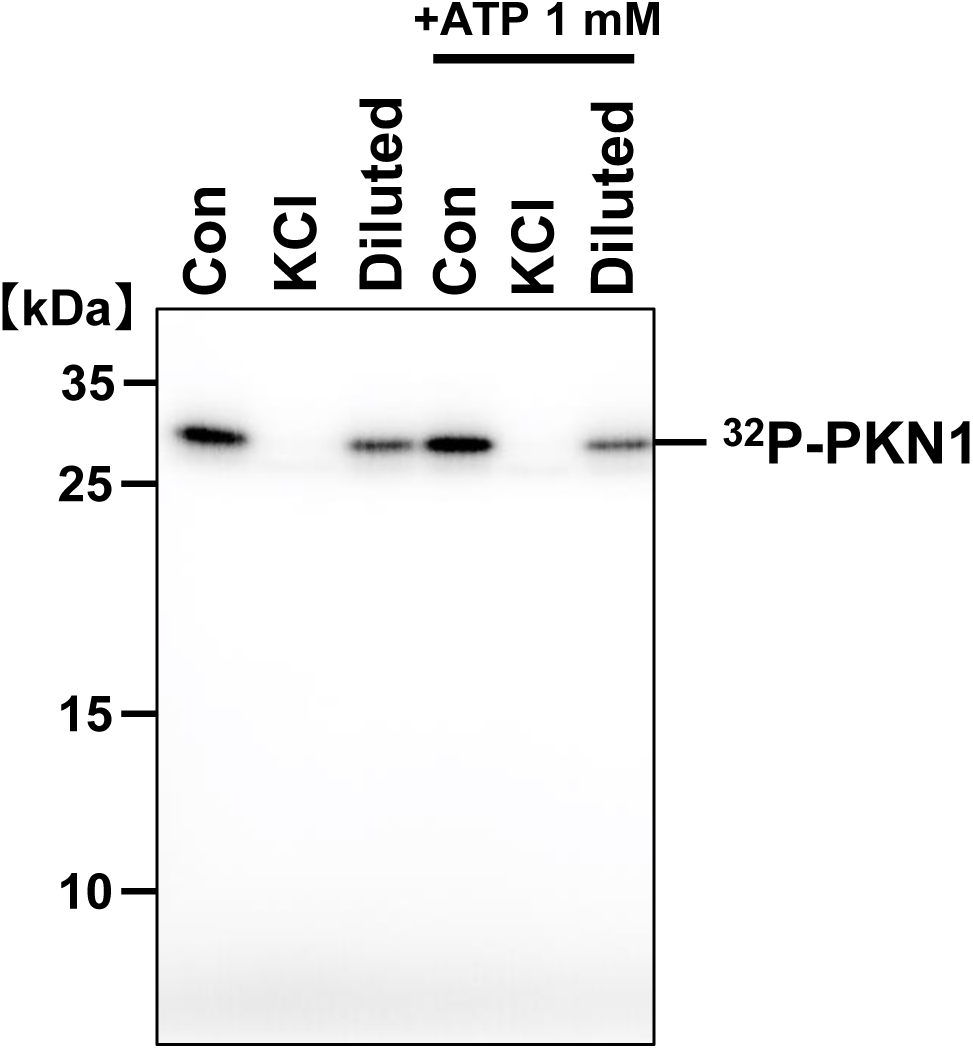
ATP-independent recovery of PKN1 activation-loop phosphorylation.COS7 extracts containing ^32^P-labeled phospho–GST–PKN1 (aa 767–788) were incubated ± 1 mM ATP with 150 mM KCl for 10 min. “Diluted” samples were generated by 1:1 mixing with 1× lysis buffer (50 mM Tris-HCl pH 7.5, 1 mM EGTA, 0.5 mM DTT, 0.1% Triton X-100) prior to autoradiography. Representative autoradiograph is shown.

**Figure 6 – figure supplement 3.**
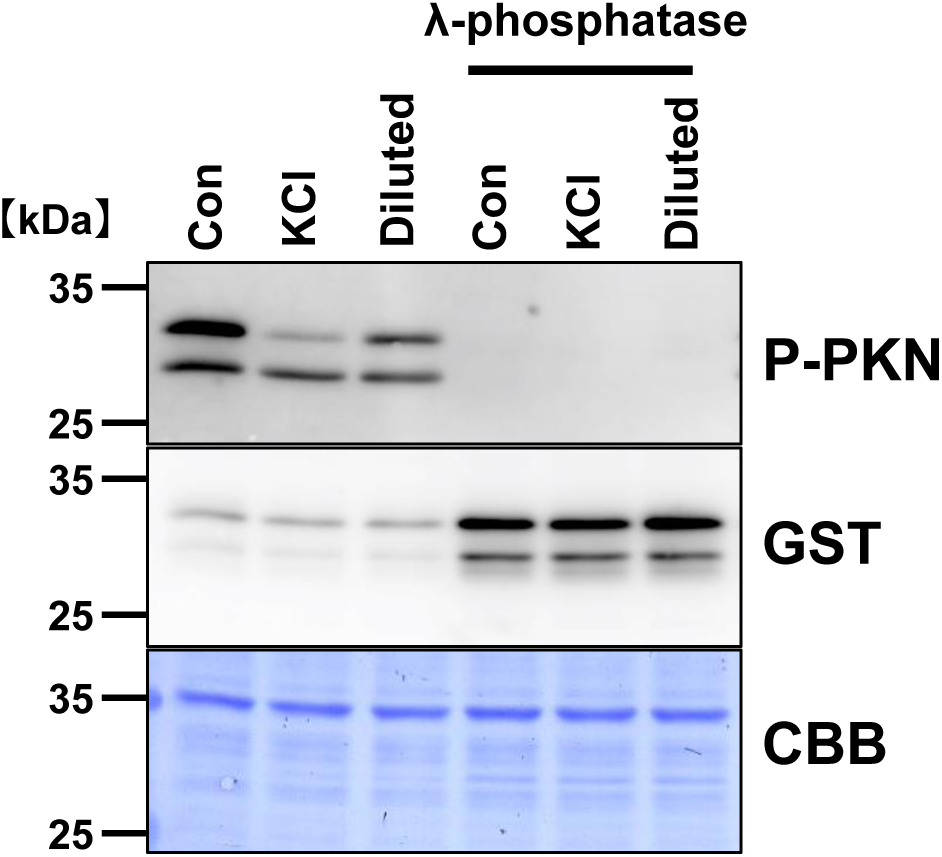
Absence of recovery with dephosphorylated PKN1 fragment. Phospho–GST–PKN1 (aa 767–788) was treated with λ-phosphatase at 30° C for 1 h, then precipitated on GST affinity beads and washed three times with 1× lysis buffer (50 mM Tris-HCl pH 7.5, 1 mM EGTA, 0.5 mM DTT, 0.1% Triton X-100) to remove residual phosphatase. COS7 extracts containing either the phospho- or λ-phosphatase–dephosphorylated fragment were incubated with 150 mM KCl for 10 min; “Diluted” samples were prepared by 1:1 mixing with 1× lysis buffer. Immunoblots for phospho-PKN, GST. Membranes were stained with Coomassie Brilliant Blue (CBB) after immunoblotting as a loading control. Representative immunoblot is shown.

